# Altered metabolic function induced by amyloid-beta oligomers and PSEN1-mutations in iPSC-derived astrocytes

**DOI:** 10.1101/2023.08.23.554346

**Authors:** R. J. Elsworthy, M. J. Finelli, S. Aqattan, C. Dunleavy, M. King, A. Ludlam, S. L. Allen, S. Prosser, R. Chen, S. Martinez Jarquin, D. H. Kim, J. Brown, H. R. Parri, S. Aldred, E. J. Hill

## Abstract

Altered energy metabolism in Alzheimer’s disease (AD) is considered a major pathological hallmark implicated in the early stages of the disease process. Astrocytes play a central role in brain homeostasis and are increasingly implicated in multiple neurodegenerative diseases. We report that astrocytes differentiated from early onset familial Alzheimer’s disease (fAD) patients or control cells treated with Amyloid β oligomers exhibit significant changes in their metabolism including glucose uptake, glutamate uptake and lactate release, with increases in oxidative and glycolytic metabolism. Furthermore, we demonstrate evidence of gliosis in fAD astrocytes in addition to a change in metabolic pathways including glutamate, purines, arginine, and the citric acid cycle. Homeostatic responses to brain activity and cellular metabolism are central to normal brain function. However, altered brain metabolism and cellular stress present significant risk factors for the onset and progression of neurodegenerative disease. This study demonstrates that fAD derived astrocytes present multiple metabolic and disease associated phenotypes early in their development suggesting that chronic alterations in fAD patient early in life that present significant risk factors for disease progression in mid-life and suggest key targets for potential diagnostic features and therapeutic agents late onset dementia in midlife.

## Introduction

Despite its relatively small proportion of total body mass (2%), the human brain requires a substantial amount of energy to sustain its complex functions, accounting for approximately 20% of total energy consumption ^1,2^ with neurons consuming the majority (75- 80%) of this energy ^3–5^. This high energy demand is attributed to the establishment of electrochemical gradients, axonal transport, and neurotransmitter recycling ^3^. Glucose is considered the primary energy substrate in the brain, although other metabolites such as ketones and lactate can also be utilized ^6,7^. Disruptions in the uptake of energy substrates by the brain can have severe consequences for normal brain function. Studies utilizing Fluorodeoxyglucose positron emission tomography (FDG-PET) have revealed a decrease in glucose uptake prior to the onset of disease symptoms in individuals at risk of developing AD, including those with familial AD (fAD), Apolipoprotein E (ApoE) 4 carriers, or patients with mild cognitive impairment ^8–11^. While neuronal loss, a hallmark of AD, could explain the decline in glucose utilization, evidence suggests that changes in energy metabolism occur before neuronal loss, potentially contributing to the development and progression of the disease^8^.

Recent evidence has increasingly implicated astrocytes, in the pathogenesis of AD. Several studies have reported altered glucose metabolism, insulin signalling, and response to oxidative stress in astrocytes in AD ^12^. ^13^. Furthermore, possession of the ApoE ε4 allele, a known risk factor for AD, is associated with significant metabolic changes in astrocytes at the earliest stages of the disease ^14–16^. These findings suggest that dysfunction of astrocytes may be an early event in AD pathogenesis, compromising the brain’s compensatory mechanisms against AD-related pathology.

Altered metabolism is recognized as an early event in the disease process in AD, understanding these changes in astrocytes is of paramount importance ^17,18^. In addition to their involvement in energy metabolism, astrocytes play crucial roles in neuronal signalling, homeostasis and neuroprotection. They form intricate interactions with neurons, creating a "tripartite synapse" and providing energetic metabolites from blood vessels to distant neurons through gap junctions ^19–21^. Astrocytes store glycogen, the brain’s exclusive glycogen reservoir, and rapidly convert it into lactate, which serves as a crucial fuel source for neurons during increased neuronal activity ^22,23^. Moreover, astrocytes exhibit neuroprotective functions by preventing glutamate toxicity and oxidative stress through the generation of antioxidants like glutathione ^24^. While numerous studies have investigated global changes in brain metabolism, redox status, gene expression, and epigenetic markers in AD, the intricate interplay between different metabolic processes, particularly in astrocytes, remains poorly understood. Brain glucose consumption is not homogeneous, and different cell types exhibit distinct glycolytic/aerobic metabolic profiles ^6^. Accumulating evidence suggests that Aβ, a key player in AD pathology, induces metabolic dysfunction in both neurons and astrocytes, possibly through mechanisms involving oxidative stress, mitochondrial dysfunction, impaired calcium signalling, and disruption of essential enzymes ^25,26^. Recent evidence of a possible role for astrocytes as the link between Aβ, tau pathology and cognitive symptoms has contributed to the increasing interest in astrocytic proteins as biomarkers of AD progression ^27,28^.

Induced pluripotent stem cell (iPSC) derived models of the AD brain are an increasingly popular tool to investigate cell type specific characteristics and cell-cell interactions. Several processes including, amyloid generation, calcium homeostasis, amino acid metabolism and neuron-astrocyte transmitophagy have been shown to be altered, in addition to displaying characteristics of a reactive phenotype ^29–32^. In this paper, the effect of Aβ oligomers on metabolic function in iPSC-derived astrocytes was investigated. We demonstrate that exposure to Aβ oligomers induces a significant shift in energy metabolism and elicits a ‘reactive’ state in healthy astrocytes. Building upon this, we characterised astrocytes differentiated from fAD patient derived iPSC cell lines carrying Presenilin 1(PSEN1) mutations. We show that PSEN1 astrocytes generate more Aβ than healthy controls and display an altered metabolic status with features of reactive gliosis. Overall, the data presented will allow for a better understanding of the underlying features of astrocyte dysfunction in AD. By focusing on the molecular mechanisms underlying astrocyte dysfunction in AD, novel therapeutic strategies may emerge, providing potential avenues to restore energy metabolism and counteract disease progression.

## Materials and methods

### Neuralisation of iPSC’s to Human Neural Precursor Cells (hNPCs)

fAD PSEN1 (R278I) iPSC line was obtained from Prof. Selina Wray (UCL, UK)^33^. The initial tissue was provided under the ethical approval of NHS Research Authority NRES Committee London-Central (REC# 08/H0718/54+5). R278I cells were cultured for expansion of cell numbers and monitored for colony size and changes in morphology before undergoing neural induction as described ^34^ based on adapted protocols previously developed ^35,36^.

### Astrocyte differentiation of Human Neural Stem Cells

Healthy control (ax0018) and PSEN1 human neural precursor cells (hNPCs) carrying L286V (ax0112) and A246E (ax0114) mutations were purchased from Axol Bioscience (Cambridge, UK) and plated as previously described ^34^ in parallel to the R278I generated hNPCs (see above). Briefly, the Control and three fAD PSEN1 cells were seeded at a density of 7×10 cells/cm in neural plating media (Axol Bioscience, Cambridge, UK) on Matrigel-matrix (356237, Corning) coated wells. After 24 hours cells were washed with D-PBS before media was exchanged for astrocyte differentiation medium (STEMdiff^tm^ Astrocyte differentiation kit #100-0013, StemCell Technologies, Cambridge, UK) and a full media exchange was completed every day for 7 days. Cells were passaged at around 80% confluency using 1 mL/well Accutase™ (A6964, Merck, UK) and dissociation was stopped with 4 mL/well ADM. Cells were plated at 5×10 /cm density and maintained in a 37°C, 5% CO2/ 95% air atmosphere with a total media exchange every other day, through two subsequent passages, before switching to astrocyte maturation media (STEMdiff^tm^ Astrocyte maturation kit #100-0016, StemCell Technologies, Cambridge, UK). Cells were seeded to a density of 2.5×10 **/**cm onto tissue culture plastic on Matrigel-matrix and media was changed every other day for 7 days or until reaching 80% confluency. Upon reaching 80% confluency and maturation, cells were passaged as before and cultured in astrocyte maintenance media (ScienCell astrocyte media, Cat #1801) until ready for analysis at day 45. Each replicate (n) was represented by a separate astrocyte induction process, from which technical repeats were generated. Cells were routinely tested for Mycoplasma contamination by luminescence assay (Lonza MycoAlert^TM^ PLUS, LT07-703). For information on cell lines used see *Table 1*.

**Table 1.** Information on cell lines used for generating astrocytes from healthy control PSEN1 fAD mutation carriers.

|  | Ax0018 | Ax0112 | Ax0113 | R278I |
| --- | --- | --- | --- | --- |
| **Diagnosis** | Healthy Control | Familial AD | Familial AD | Familial AD |
| **Sample type** | Dermal Fibroblast | Dermal Fibroblast | Dermal Fibroblast | Dermal Fibroblast |
| **Donor sex** | Male | Female | Female | Male |
| **Age at sampling (yrs)** | 74 | 38 | 31 | 60 |
| **Age of onset (yrs)** | n/a | 39 | 45 | 58 |
| **Karyotype** | Normal | Normal |  |  |
| **Reprogramming method** | Episomal Vector | Episomal Vector | Episomal Vector | Episomal Vector |
| **Induction method** | Monolayer – Axolbio NIM<sup>35</sup> | Monolayer – Axolbio NIM<sup>35</sup> | Monolayer – Axolbio NIM<sup>35</sup> | Monolayer – NIM<sup>35,79</sup> |
| **Mutation** | None | PSEN1 (L286V) | PSEN1 (A246E) | PSEN1 (R278I) |
| **APOE status** | ε2/ ε2 | ε3/ ε3 | ε2/ ε3 | ε2/ ε4 |

Note: Adapted from<sup>34</sup>

**Table 2.** Neural induction media (NIM) made by the addition of small molecule signalling pathway inhibitors to Essential 6 as a base medium.

| COMPONENT | AMOUNT |
| --- | --- |
| GIBCO™ ESSENTIAL 6™ MEDIUM (THERMOFISHER SCIENTIFIC, A1516401) | 50ml |
| XAV939 (STEM CELL TECHNOLOGIES, 72672) | 10μl |
| LDN193189 (STEM CELL TECHNOLOGIES, 72147) | 0.5μl |
| SB431542 (STEM CELL TECHNOLOGIES, 72232) | 50μl |

Note: from<sup>34</sup>

### Immunocytochemistry

Cells were fixed in 4% (v/v) paraformaldehyde (PFA) in D-PBS. The cells were then incubated for 10 minutes in PBS with 0.2% (v/v) Triton X-100 followed by blocking for 1hr in PBS containing 0.2% (v/v) Triton X-100 and 3% (w/v) bovine serum albumin (A9418 Sigma-Aldrich, UK). Primary antibodies for ALDH1A1 (PA5-32127, Invitrogen), Glial fibrillary acidic protein (14-9892-82, Invitrogen) and S100β (PA5-78161, Invitrogen) were diluted in blocking buffer and added for 1hr. Following primary antibody incubation, cells were washed with blocking buffer and appropriate secondary antibodies, Alexa Fluor® 488 AffiniPure Goat Anti-Rabbit IgG (1:2000, 111-545-144, Jackson Laboratories) and Alexa Fluor® 633 Goat anti-Mouse IgG (1:2000, A-21052, ThermoFisher Scientific) were added for 1hr. Cells were mounted in Prolong^TM^ Gold Antifade Mountant with DAPI (P3935, ThermoFisher Scientific) to glass slides and imaged using an EVOS m5000 imaging system (AMF5000, ThermoFisher, UK).

### Preparation and treatments of Synthetic Aβ1-42 Oligomers

Human HFIP Aβ1-42 (AG968, Sigma Aldrich) was prepared in oligomeric form as previously described ^37^. Briefly, Human HFIP Aβ1-42 was resuspended in DMSO to 5mM. Monomers were diluted in F-12 culture media, without phenol red, to a concentration of 100μM and incubated for 24hrs *at 4°C.* To determine the possible toxic effects of human Aβ1-42 oligomers on the metabolism of astrocytic cells derived from ‘healthy’ patient hNPCs, cells were treated with oligomeric Aβ1-42 over a range of concentrations based on our previous studies ^38^. Astrocytes were incubated for either 4hrs or 48hrs at 37°C in a humidified atmosphere of 5% CO2. Subsequently, conditioned media (CM) was collected and centrifuged at speed of 200 x g for 5 minutes. Protein lysates were collected from cells using ice cold RIPA buffer (R2078, Sigma Aldrich) with Halt^TM^ 100x protease inhibitor cocktail added (78440, Thermofisher). CM and lysates were transferred into 1.5ml sterile microcentrifuge and stored at -80°C for future measurements.

### Total Cellular Protein Quantification

The protein concentration of cell lysates was determined using a modified protocol of the Bicinchoninic acid (BCA) protein assay ^39^ to enable standardisation of assays.

### MTT assay

iPSC derived astrocytes were seeded into 96 well plates at a density of 8×10^5^ per well. Triplicate technical repeats for each control and experimental condition were used. For the assay, 3-(4,5-dimethylthiazol-2-Yl)-Diphenyltetrazolium Bromide (MTT, CT01-5, Sigma-Aldrich, UK) stock solution was diluted in F12 medium without phenol red (1:5), added to each well and incubated for 3hrs (37°C). The MTT solution was then aspirated and DMSO (50μl) was added to each well. Cells were placed on a plate shaker (500rpm) for 30 seconds followed by incubation for 10 minutes (37°C). Finally, absorbance was read at 590nm (Fluostar Omega, BMG Labtech).

### AOPI cell viability count

The viability of astrocytes following treatment with Aβ1-42 oligomers was measured using an automated dual fluorescence cell count (Cellometer 2000, Nexcelom). Briefly, astrocytes were treated with either 0.2µM,1µM or 2µM Aβ1-42 and compared to the untreated control for 48hrs. Acridine orange (AO) and Propidium iodide (PI) fluorescence imaging was quantified to determine live/dead, cell size and total cell counts (ViaStain™ AOPI Staining Solution, Nexcelom).

### Seahorse analytics

Oxygen consumption rate (OCR), extracellular acidification rate (ECAR), mitochondrial respiration indicators and glycolysis indicators, respectively, were measured from live cells in real time using Seahorse Extracellular Flux (XF) 24 Analyzer (Agilent Technology). OCR and ECAR were measured simultaneously. Matured astrocytes were plated on Matrigel-coated XF24 TC plate at a density of 40,000 cells/well in astrocyte maturation medium and cultured overnight. For Aβ treatment comparisons, 0, 0.2, 1 & 2μM were added to control cells 48hrs prior to commencing the assay. Before conducting the assay, media was changed to Seahorse XF Assay medium (Seahorse RPMI supplemented with 2 mM Glutamine, 1mM Sodium pyruvate and 10mM glucose). Oligomycin (2μg/mL), BAM-15 (3μM), antimycin/rotenone (2μM) and 2-Deoxyglucose (50mM) injections were performed in line with manufacturer’s instructions using 3 min mix time, 3 min wait time and 3 min measure cycle. Following the assay, results were normalised using CyQUANT nucleic acid stain (C7026, Invitrogen). Data was analysed using Wave program (Agilent Technology).

### Glycogen assay

Astrocytes cultured in 12 well plates were scraped into 300uL ice cold HCL (30mM), sonicated for 15 seconds before dilution 1:2 with D-PBS, and the diluted samples were mixed with 0.1M acetate buffer at a pH of 4.6. The diluted sample was divided equally into two separate microfuge tubes, one tube to measure the amount of glycogen and the second tube to measure the free glucose. Next, an equal volume 1mg/ml Amyloglucosidase enzyme stock (AMG.S) (Sigma-Aldrich, UK), which was prepared by adding 75µl of Amyloglucosidase enzyme reagent (AMG) in 1ml of acetate buffer with a pH of 4.6. Afterwards, all the samples were incubated at 57.5°C on a heat block for 2hrs. After incubation, 30µl of sample was transferred into a 96-well plate and added 100µl of Hexokinase enzyme reagent (HK) (Sigma-Aldrich, UK), and then mixed and incubated at room temperature for 15mins (Fig. 2.3). The absorbance was read at 370nm using Thermo multiscan EX 96-well plate reader (Thermofisher, UK). The protein content levels in the cell lysate samples were determined using BCA assay in order to normalise the glycogen values. The ability of iPSC derived astrocytic cells to store, and breakdown glycogen was assessed using physiological cues such as hypoglycaemia: 1,4-dideoxy-1,4-imino-d-arabinitol (DAB) (Sigma-Aldrich, UK) or drugs including: Dibutyryl cyclic adenosine monophosphate (dbcAMP) (Tocris, UK), Isoproterenol (Tocris, UK), Ouabain (Tocris, UK) and DL-threob-benzyloxyaspartic acid (TBOA) (Tocris, UK).

### Glucose assay

Glucose levels in culture media were quantified using a bioluminescent NADH detection assay according to manufacturer’s instructions (Glucose-glo, Promega). Briefly, the addition of glucose detection reagent 1:1 to samples (12.5µL) in a 384-well plate was incubated for 1 hour or until a stable luminescent signal was achieved. Luminescence was plotted against serially diluted standards from 50-0.8µM to calculate glucose levels in cell conditioned culture media.

### Lactate and glutamate assays

Glutamate (ab83389, Abcam) and Lactate (65331, Abcam) levels were quantified in cell conditioned media by colorimetric assays as per the manufacturer’s instructions. For glutamate uptake and subsequent lactate release, a modified protocol from Mahmoud et al. (2019) was utilised. Briefly, astrocytes were seeded in 12-well plates at 3.5 x 10^4^ and cultured for 24hrs before treatment with Aβ for 48hrs. Cells were washed 2 times with Hank’s Balanced Salt Solution (HBSS, Gibco™ 14025092) containing Ca^2+^ before incubation for 4hrs (37 °C, 5% CO2) with 200µM glutamate. Glutamate uptake by astrocytes was measured by subtracting the amount of glutamate measured in the CM from the amount added to the cells. The protein content levels in the cell lysate samples were also determined using BCA assay to normalise values ^40^.

### Immunoassays

Immunoassays for Aβ1-40 (ThermoFisher, KHB3481), Aβ1-42 (ThermoFisher KHB3441), aggregated Aβ (ThermoFisher, KHB3491) and sAβPP-α (MyBioSource, MBS9358454) in CM was measured via ELISA according to manufacturer’s instruction. Media samples were first concentrated using Amicon^®^ ultra-15 centrifuge filter units (3kDa UFC9003, Millipore). IL-6 and IL-8 levels in conditioned media were quantified using Quantikine immunoassays (HS600C and D8000C, R&D systems). GFAP was measured in CM using Human GFAP DuoSet ELISA (R&Dsystems, DY2594-05) as per manufacturer’s instructions.

### Flow cytometry

To quantify intracellular accumulation of IL-6 and IL-8, astrocytes (1×10^6^) were incubated with protein transport inhibitor containing Brefeldin A (555029 Golgiplug, BD Biosciences) and Monensin (554724 Golgistop, BD Biosciences) following 48hrs treatment with Aβ oligomers or 4hrs with IL1β (10ng/mL, 200-01B Peprotech). Astrocytes were then dissociated and washed two times with D-PBS before fixing and permeabilisation for 20 minutes (554714 Cytofix/Cytopearm, BD Biosciences). Astrocytes were washed a further two times in wash buffer before incubation with IL-6-PE (12-7069-82, Invitrogen) and IL-8 FITC (BH0814, BioLegend) conjugated antibodies for 30 minutes. Following a final three washes, astrocytes were analysed using a flow cytometer. Fluorescent compensation was applied using Ultracomp ebeads (01-2222-42, Invitrogen) and unstained astrocytes.

### ADAM10 activity

ADAM10 activity was measured via fluorometric FRET assay following manufacturer’s instructions (AS-72226, Sensolyte 520, Anaspec) and as previously described^41^. ADAM10 activity was calculated using linear regression in relative fluorescence units (RFU) compared to 5-FAM peptide activity.

### Oxidative stress measurements

Superoxide generation via MitoSOX^TM^ Red mitochondrial superoxide indicator (M36008, Invitrogen) and lipid peroxidation via total F2 isoprostanes (8-isoprostane ELISA kit, Cayman Chemical) were quantified as previously described ^41^. Protein Carbonylation was assessed by the method of Carty *et al*. (2000). Briefly, cell lysates and standards (BSA) were added to carbonate buffer (sodium carbonate 50mM, pH 9.2) and plated into 96 well plates (50µl at 0.05mg/ml) in triplicate. Proteins were allowed to bind for 1hr at 37°C before washing with TBS–Tween (0.5%). DNPH was added in HCl (1mM) and allowed to react for 1hr at room temperature before washing with TBS–Tween (0.5%). Non-specific binding sites were blocked overnight at 4°C with TBS-Tween (1%). After washing, rabbit anti-DNPH primary antibody (1:1000) was applied and incubated for 1hr at 37°C and, following washing with TBS-Tween (1%), anti-rabbit IgE conjugated to peroxidase (1:5000) was also incubated at 37°C for 1hr. The reaction was visualised by o-phenylenediamine tablets with hydrogen peroxide (final conc. 7.8mM) in citrate-phosphate buffer (10mL) and stopped by addition of sulphuric acid (2N). Absorbance was read at 490nm (Fluostar Omega, BMG Labtech).

### Metabolites extraction and analysis

Metabolite extraction and profiling were performed as previously described ^42^. Briefly, at day 50 in culture, cells (5×10^5^) were seeded in 6 well plates and maintained for 48hrs before a full media exchange was completed. Following a further 48hrs, 1 ml of the conditioned medium in each culture, was collected and centrifuged at 200 x g for 5 mins. 250ul of the collected CM were transferred to a new microcentrifuge tube for extraction and protein precipitation. 750ul of ice-cold methanol was added and vortexed for 1min and incubated at - 20°C for 20 min to precipitate proteins. Samples were then vortexed again for 15sec and centrifuged at 17,000 rpm for 10mins at 4℃. The supernatant was transferred into precooled fresh tubes and samples were stored at -80°C until LC-MS analysis. Metabolite extraction from control, mutant astrocytes and Aβ-treated control astrocytes were prepared and processed in parallel, as well as no cells controls (blanks). A pooled quality control (QC) sample was prepared by mixing equal volume from each sample (excluding blanks) to evaluate the robustness, performance and stability of the analytical system. Experimental and blank samples were randomised before injection. QC samples were included throughout the analysis to check the performance of the analytical system. Five mixed authentic standards solutions (total of 268 standards) were injected under the same condition for metabolite identification.

LC-MS-based metabolite profiling was performed on an Dionex Ultimate 3000 HPLC system coupled to a Q-Exactive Plus hybrid quadrupole-Orbitrap mass spectrometer (Thermo Fisher Scientific, UK) as previously described ^43^. Samples were separated on a ZIC-pHILIC column (5μm, 4.6´150mm) from Merck Sequant (Watford, UK), with the mobile phases of 20 mM ammonium carbonate (A) and acetonitrile (B). Starting with 20%, mobile phase A increased linearly to 95% within 15 min before decreasing back to 20% in 2 min, where it was held for 7 min for equilibration. The injection volume was 10μL and the flow rate was 300 μL/min. The chamber of autosampler was kept at 4 ℃ and column temperature was maintained at 45℃. Mass spectrometer, fitted with HESI source, rapidly switched between positive (ESI+) and negative (ESI-) mode with the spray voltage +4.5 kV and -3.5 kV, respectively. Other settings were optimized as follows: capillary temperature, 275℃; sheath gas flow rate, 40 arb; aux gas flow rate, 5 arb; sweep gas flow rate, 1 arb; S-lens RF level, 55%. The samples were acquired using full MS scan method, ranging m/z 70∼1050 at 70,000 resolution. Top 10 data-dependent MS/MS (ddMS/MS) was performed on QC samples at resolution of 17500 and stepped normalized collision energy of 20, 30 and 40.

LC-MS data was analysed with Compound Discoverer 3.3 SP1with an untargeted metabolomic workflow (Thermo Fisher Scientific, UK) for peaks picking, peaks alignment, gap filling and metabolite identification. Compound annotation was made using exact mass (5 ppm error) using Human Metabolome Database (HMDB), Kyoto Encyclopedia of Genes and Genomes (KEGG), the accurate mass and retention time of authentic standards, and *mz*Cloud fragmentation database.

According to the metabolomics standards initiative and scale ^44^, the metabolites with high identification confidence (Level 1 and Level 2) were included and reported. In this study, Level 1 identification was that metabolites were matched with accurate masses, retention times and MS/MS fragmentation of authentic standards. Level 2 identification was detected peaks matched with accurate masses and MS/MS information of compounds in spectral library when lack of standards.

All identified compounds were subjected to pathways analysis using the online platform of MetaboAnalyst 5.0. Non-human putative metabolites were excluded. The abundances of metabolites were log-transformed, and Pareto scaled before mapping the KEGG pathways and generating networks of interacting biological entities.

### Statistical analysis

All quantitative data in the text and figures are presented as Mean ± S.D. unless otherwise stated. To generate a sigmoidal dose-response curve for Aβ treatment or standard curves from plate-based assays either linear regression or 4-parameter logistic regression was used to plot known concentrations against optical absorbance at specified wavelengths. From this sample concentrations were calculated and normalized to total protein concentration in corresponding cell lysate. Significance was calculated using individual t-tests for grouped data or using ordinary one-way ANOVA with Bonferroni *post hoc* corrections and using linear regression models. All data was processed using GraphPad Prism (Version 9.3.1).

For metabolomics analysis, data was log-transformed, and student t-test p value was adjusted with false discovery rate using Benjamini-Hochberg approach, compensating for the multiple testing problem. Adjusted *p* < 0.05 was regarded significant. Multivariate analysis, including principal component analysis (PCA) and orthogonal partial least squares discriminant analysis (OPLS-DA) was performed on SIMCA-P v16 (Umetrics, Sweden) to determine significant metabolites between each sample groups. After log transformation, the method of UV scaling and Pareto Scaling were used for PCA and OPLS-DA respectively to make data more reasonable and comparable.

## Results

### iPSC-derived astrocytes display functional metabolic glycogen mobilisation in response to hypoglycaemia and adrenergic receptor activation treatment

To confirm differentiation of astrocytes from hNPCs, cellular changes in morphology was monitored via phase-contrast imaging. Astrocytes were identified and stained using ICC for the astrocytic markers S100β, GFAP and ALDH1 at DAY 45+ (Figure 1-A).

**Figure 1:**
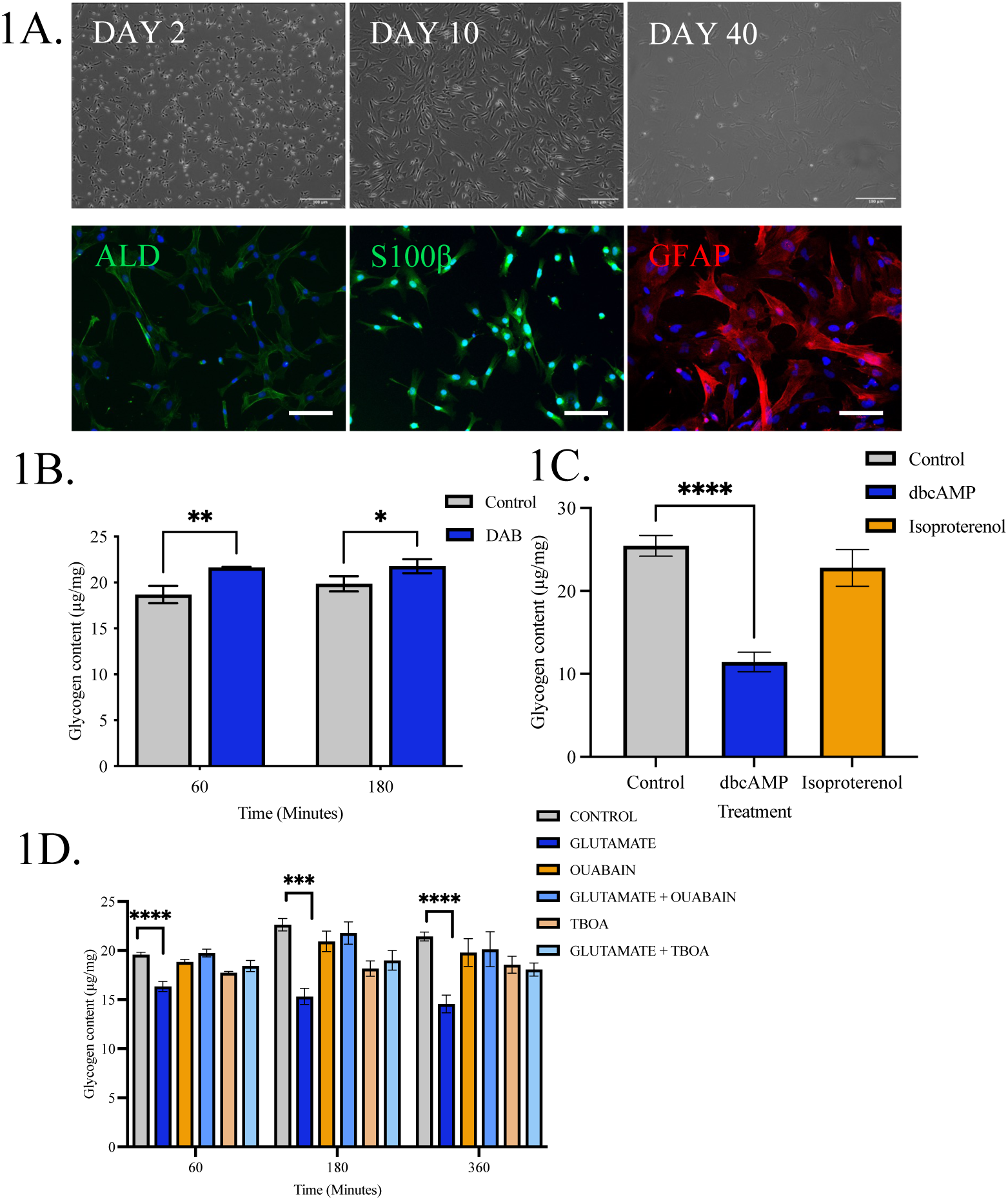
Characterisation of ‘Healthy’ iPSC derived Astrocytes. A) Representative images showing differentiation and ICC staining of iPSC derived astrocytic cells at day 45+. The astrocytes were differentiated from ‘healthy’ control patient NPCs for >40 days using astrocytes differentiation and maturation protocols. TOP) Phase contrast images of control astrocytes on days 2, 10 and 40 in culture post differentiation. Bottom) Cells were stained using immunofluorescent antibodies for astrocytic markers ALDH1A1 (Left, green), S100β (Middle, Green) and GFAP (Right, red), nuclei were counterstained in each image with DAPI (blue). Scale bars: 100μM. 1B) Cellular glycogen content of control astrocytes following exposure to hypoglycaemic conditions and treatment with DAB over 60 and 120 minutes. 1C) Glycogen content of cells treated with dbcAMP an isoproterenol. Glycogen contents of cells treated with Glutamate, oubain, glutamate and Ouabain, TBOA and Glutamate and TBOA for 60, 180, and 360 minutes. Results are expressed as ug/mg protein ± SD, n=3 P<0.05 (*), P<0.01 (**), P<0.001 (***). For DAB (1B) a two-way analysis of variance (ANOVA) was performed follow by Sidaks post-test. Comparisons between treatments (1C) were performed using analysis of variance (ANOVA) followed by Dunnet’s post-test.

To determine whether functional glycogen stores were present in iPSC-astrocytes, control cells were cultured under glucose-starvation conditions to induce intracellular glycogen breakdown and with inhibitors of glycogen breakdown. During this period, glycogen content of cells was measured. Glycogen breakdown was significantly blunted with differences in cellular glycogen content from 18.69 ± 0.55µg/mg in the control at 60min (*p*<0.001) to 21.64 ± 0.03µg/mg in the presence of DAB (glycogen phosphorylase and synthase inhibitor) and from 19.86 ± 0.47µg/mg in the control at 180 minutes to 21.78 ± 0.40 µg/mg (*p*<0.01) (Figure 1-B). Next, to stimulate glycogen breakdown, control astrocytes were incubated with Isoproterenol (a β1 and β2 adrenergic agonist) or dbcAMP (an inducer of glycogen phosphorylase activity) for 180 minutes (to allow time for glycogen breakdown). Cellular glycogen content was assessed after 180 minutes. dbcAMP induced significant reduction in glycogen content from 25.44 ± 1.25µg/mg, to 11.44 ± 1.17µg/mg (*p*<0.0001), however there was no significant change in glycogen storage following treatment with Isoproterenol (Figure 1-C).

Given the key role astrocytes play in the homeostasis of the neurotransmitter glutamate levels and the energy cost associated with glutamate uptake, the effect of glutamate on glycogenolysis was assessed. Astrocytes were treated with glutamate (1mM) with and without Ouabain (100µM, an inhibitor of Na^+^/K^+^ ATPase) and TBOA (100µM, glutamate transporters inhibitor). Intracellular glycogen levels were measured at 60, 180 and 360 minutes corresponding to the timescale of glycogen breakdown to occur in astrocytes ^45^. Glutamate treatment induced significant reduction in glycogen levels in control astrocytes at 60min (19.59 ± 0.24µg/mg vs. 16.34± 0.51µg/mg, *p*<0.0001), 180min (22.63 ± 0.63µg/mg vs. 15.32 ± 0.82µg/mg, *p*<0.0001) and 360min (21.43 ± 0.45µg/mg vs. 14.56 ± 0.91µg/mg, *p*<0.001). Treatment with Ouabain or TBOA with and without glutamate blocked the effect of glutamate on glycogenolysis (*p*>0.05) (Figure 1-D). The results are consistent with the physiological situation where glutamate uptake by EAATs is dependent on the electrochemical gradient maintained by the Na/K ATPase, and additionally that astrocyte glycogen can be a source for ATP.

### Amyloid-beta oligomer treatment induces metabolic alterations and impaired glutamate uptake in iPSC-derived astrocytes

Next, we investigated whether stressors observed in AD, such as Aβ, impact astrocyte glycolytic metabolism. Here we focused on treatment with Aβ1-42 oligomers as a model for acute exposure to Aβ in AD. First, we determined whether Aβ1-42 oligomers affected cellular metabolic enzyme activity using MTT and viability of control astrocytes with AOPI probes. A dose-dependent response showed that 4hr treatment with Aβ1-42 oligomers (0.078-5µM) caused a significant reduction in metabolic enzyme activity (78.77 ± 6.57%, *p*<0.02) at 0.625µM compared to control. This was with a maximum activity of 104.60%, minimum activity of 66.41% and EC50 of 0.571µM. After 48hr treatment with Aβ1-42 oligomers (0.078- 5µM), there was a significant reduction in metabolic enzyme activity (79.33 ± 7.13%, *p*<0.01) at 0.078µM compared to control. This was with a maximum activity of 99.31%, minimum activity of 60.80% and EC50 of 0.099µM. As MTT viability is reliant on mitochondrial function and enzymatic reduction of MTT, we also quantified viability using AOPI dual fluorescence imaging which is an indication of cell membrane permeability. Treatment of astrocytes with Aβ1-42 oligomers significantly reduced cell viability at 0.2µM (88.77 ± 1.468 %, *p* = 0.014), 1µM (88.77 ± 0.986 % *p* = 0.014) and 2µM (85.18 ± 0.745 % *p* < 0.001) compared to control (92.20 ± 1.158 %).

Based on our previous functional data, astrocytic glycogen levels were determined in cell lysates after 48-hour treatment with Aβ1-42. There was a significant reduction in glycogen content following exposure with 2µM Aβ1-42 oligomers (12.36 ± 1.94µg/mg, *P<0.01*) compared to control (28.97 ± 3.23µg/mg). No significant effects were observed following treatment with either 0.2µM or 1µM Aβ1-42 (Figure 2-D). To determine the effect of Aβ1-42 oligomers on glucose uptake in astrocytes, the amount of glucose remaining in the media was monitored. Following treatment with Aβ1-42 there was a significant increase in glucose remaining in culture media after 48hrs following 2µM treatment (15990 ± 3731µM/mg, *P<*0.035) compared to control (6644 ± 2756 µM/mg) (Figure 2-E). No significant difference in glucose levels in the media (*p*>0.05) were detected at any other time point or concentration of Aβ1-42 treatment.

**Figure 2.**
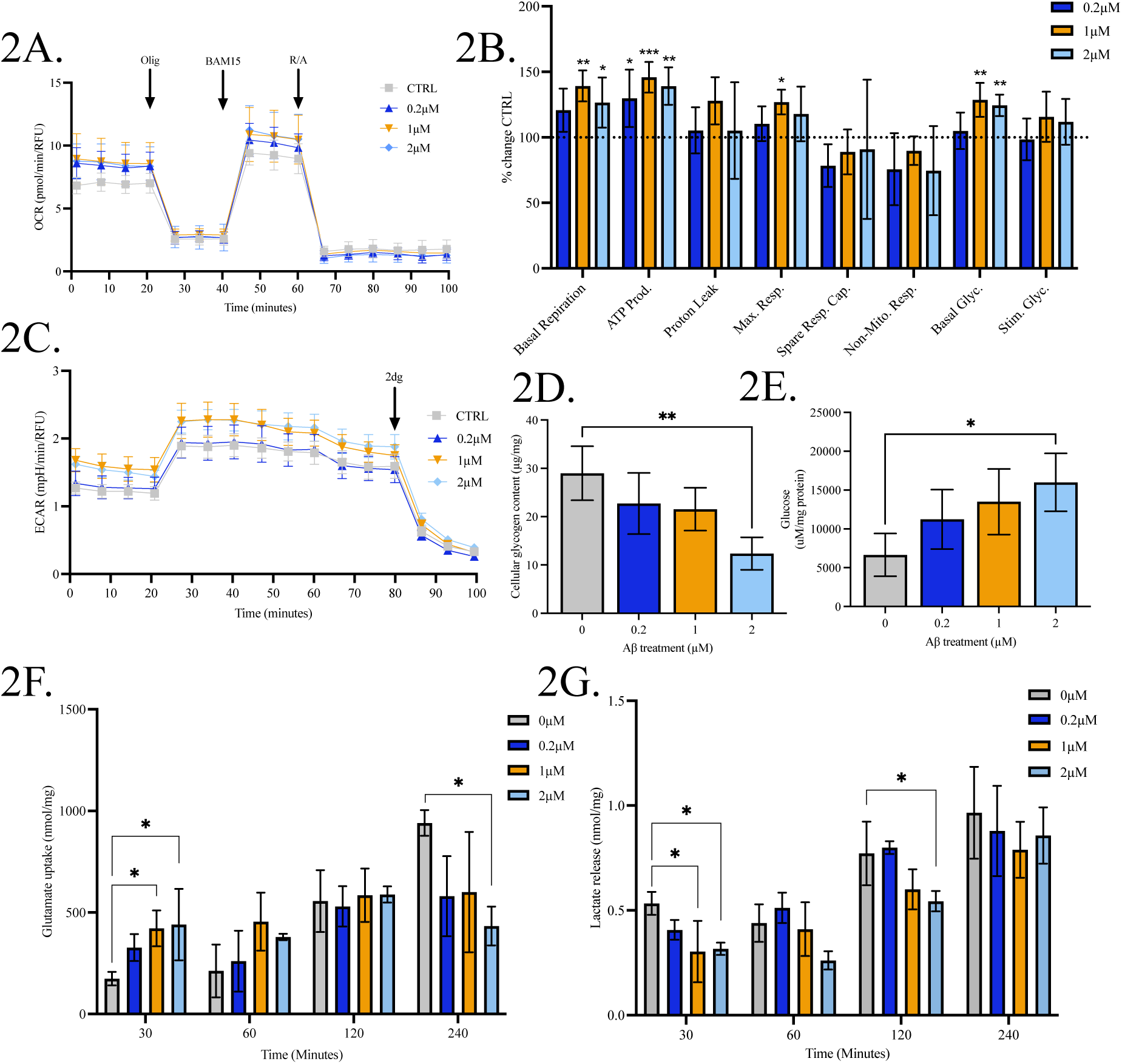
Amyloid β treated control astrocytes exhibit difference in their bioenergetic profiles and metabolite processing. Astrocytes (45 days old) were treated with 0.2, 1, 2µM Aβ oligomers. A) Profiles of Seahorse XFp Mito Stress Test data for oxidative consumption rates (OCR, pmol/min) C) Extracellular acidification rate (ECAR, mpH/min) measured after treatment with 0.2, 1, 2µM Aβ oligomers. B) Percentage change of treated cells over control (untreated) cells. D) Cellular glycogen content of cells (µg/mg cellular protein). E) Glucose levels remaining in the media µM/mg cellular protein, F) Glutamate uptake (nmol/mg cellular protein), Lactate release (nmol/mg cellular protein) after treatment with 0.2, 1, 2µM Aβ oligomers. Results are expressed as ± SD, n=3 P<0.05 (*), P<0.01 (**), P<0.001 (***). Comparisons between treatments were performed using ANOVA followed by Dunnet’s post-test.

To understand whether discrete cellular bioenergetic profiles were altered by Aβ1-42 oligomers, metabolic activity was analysed in the astrocytes using Seahorse XF analyser to gain real-time data for both oxygen consumption rate (OCR) and extracellular acidification rate (ECAR). Basal OCR was significantly increased following treatment with either 1µM (9.691 ± 0.824 pmol/min, *p* = 0.001) or 2µM (8.810 ± 1.33 pmol/min, *p* = 0.03) Aβ1-42 compared to control (8.404 ± 1.145 pmol/min). Maximal respiratory capacity was significantly increased only at treatment with 1µM Aβ1-42 (11.68 ± 0.865 pmol/min, *p* = 0.025) as compared to vehicle-treated (9.203 ± 1.102 pmol/min). ATP-coupled respiration was significantly increased at 0.2µM (5.691 ± 0.960 pmol/min, *p* = 0.018), 1µM (6.398 ± 0.513 pmol/min, *p* < 0.001) and 2µM (6.101 ± 0.626 pmol/min, *p* = 0.002) Aβ1-42 treatments compared to control (4.386 ± 0.429 pmol/min). There was no significant difference in proton leak, spare respiratory capacity, or non-mitochondrial oxygen consumption (Figure 2-A&B).

For ECAR measurements, basal glycolysis was significantly elevated following treatment with both 1µM (1.575 ± 0.584 mpH/min, *p* = 0.002) and 2µM (1.524 ± 0.100 mpH/min, *p* = 0.008) Aβ1-42 compared to control (1.225 ± 0.098 mpH/min). There was no significant difference in glycolytic rates between conditions following oligomycin injection. Glycolysis derived ATP levels were significantly elevated following 1µM (2.243 ± 0.223 pmol/min, *p* = 0.017) and 2µM (2.231 ± 0.149 pmol/min, *p* = 0.002) Aβ1-42 treatments compared to control (1.860 ± 0.134 pmol/min, *p* = 0.021) (Figure 2-C).

Our data has demonstrated that astrocytes increase glycolysis and glycogenolysis after stimulation with glutamate in order to meet increased energetic demands. Given that utilisation of alternative energy sources may become essential in AD wherein levels of lactate become elevated in MCI and CSF of AD patients and glutamate levels may become perturbed, we next investigated the impact of Aβ ^46^. To understand whether the effect of Aβ1-42 treatments was associated with alterations in glutamate uptake and lactate release, astrocytes were treated with 200mM glutamate and sampled at 30 minutes, 60 minutes, 120 minutes, 4 hours. Glutamate and lactate levels in conditioned media was also collected after 48hrs. After 30 minutes, glutamate uptake was significantly elevated following 48-hour treatment with 1µM (421.7 ± 88.41 µmol/mg, *p* = 0.004) and 2µM (440.3 ± 175.8 µmol/mg, *p* = 0.031) Aβ1-42 compared to control (164.1 ± 37.64 µmol/mg). There was no significant difference between glutamate uptake at both 60 and 120 minutes. Interestingly, by 4hrs glutamate uptake was reduced following Aβ1-42 treatments and was significantly lower than control (940.7 ± 63.25 µmol/mg) following 2µM treatment (433.2 ± 95.57 µmol/mg, *p* = 0.026). Glutamate levels in CM were not significantly altered after 48hrs (Figure 2-F). Lactate release was significantly lower after 30-minute treatment with 200mM glutamate after both 1µM (0.304 ± 0.146 nmol/mg, *p* = 0.024) and 2µM (0.317 ± 0.029 nmol/mg, *p* = 0.031) Aβ1-42 treatments compared to control (0.533 ± 0.055 nmol/mg). After 60 and 120 minutes, lactate release remained lowered, however this was only significant at 120 minutes following 2µM Aβ1-42 treatment (0.544 ± 0.049 nmol/mg, *p* = 0.044) compared to control (0.772 ± 0.152 nmol/mg). There was no difference in lactate release after 4 or 48hrs (Figure 2-G).

### iPSC-derived Astrocytes exposed to amyloid oligomers display features of reactive gliosis

As changes in metabolism are well-known to drive changes in astrocyte phenotype from neuroprotective to neurotoxic ^47^, we next investigated whether Aβ1-42 treatment induced gliosis in astrocytes. One key feature of reactive gliosis is an increase in cell size ^48^. Following treatment with Aβ1-42 a detectable increase in cell diameter at 0.2µM (10.57 ± 0.351 microns, *p* = 0.018), 1µM (10.67 ± 0.635 microns, *p* = 0.012) and 2µM (10.90 ± 0.265 microns, *p* = 0.004) compared to control (9.433 ± 0.058). Further evidence of an altered immune state was gathered by quantifying the accumulation of two cytotoxic cytokines secreted by reactive astrocytes, IL-6 and IL-8, was quantified by ELISA following treatment with IL1-β as compared with vehicle-treated condition. IL1-β treatment significantly increased both intracellular IL-6 (6.865 ± 1.569 FC, *p* = 0.001) and IL-8 (3.360 ± 0.530 FC *p* < 0.001) in comparison to control (values expressed fold change to control). In response to Aβ1-42 treatment, IL-6 (4.080 ± 1.019 FC, *p* = 0.02) was significantly elevated, but this was not seen for IL-8 (1.618 ± 0.113 FC, *p* > 0.05) compared to control. Interestingly, after 48hrs secretion of in IL-6 (32.09 ± 3.642 pg/mg, *p* = 0.04) and IL-8 (17.89 ± 3.240 pg/mg, *p* = 0.03) were significantly increased as compared to control (15.27 ± 2.877 pg/mg and 11.90 ± 1.410 pg/mg respectively) following IL1-β treatment. With 48hr treatment of 2µM Aβ1-42, astrocytes showed a significant increase in IL-6 (27.76 ± 6.871 pg/mg, *p* = 0.02) but not for IL-8 (15.10 ± 1.636 pg/mg, *p* > 0.05) (Figure 3-A-D). In addition to cytokine markers, GFAP levels was also quantified in both media and cell lysates using an ELISA. GFAP is increasingly recognised as a marker of neurodegenerative disorders as well as traumatic brain injury and is upregulated during gliosis in response to chemical, biological insults as well as brain trauma ^49^. There was no significant difference in secreted GFAP in CM following 48hr of Aβ treatment, however, 2µM treatment with Aβ significantly elevated intracellular GFAP protein levels (20.26 ± 0.952 ng/mg) compared to control (18.25 ± 0.851 ng/mg). Quantification of mitochondrial superoxide generation relative to control (fold change, FC) showed a dose depended lowering following treatment with 2µM Aβ1-42 (0.837 ± 0.054 FC, *p* = 0.015) (Figure 3-E&F).

**Figure 3.**
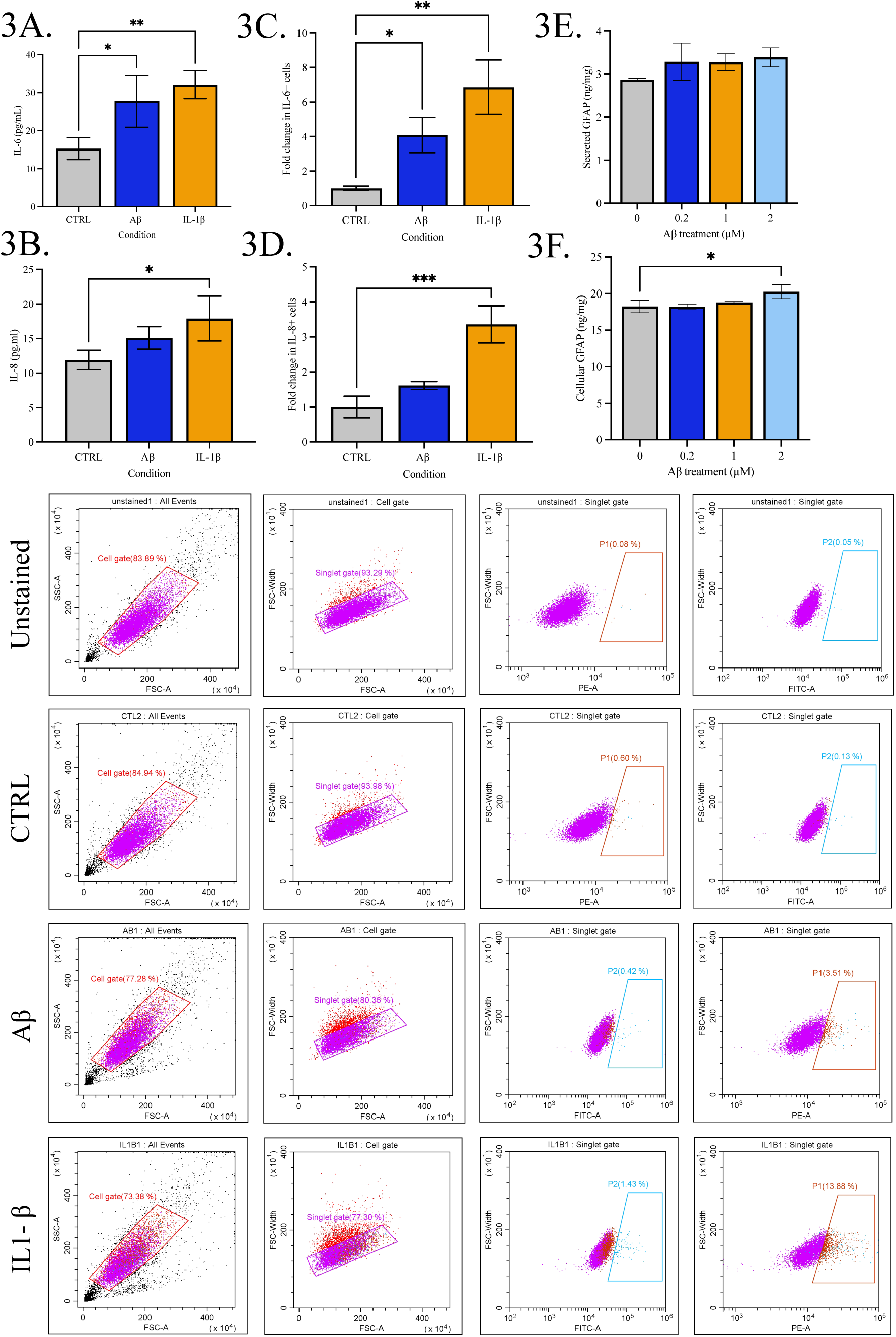
Astrocytes display markers of gliosis following exposure to Aβ oligomers. Control astrocytes were treated with Aβ oligomers for 48 hours. Astrocytes (45 days old) were treated with 0.2, 1, 2µM Aβ oligomers. Cytokine or GFAP levels in media were measure using ELISA or by investigating intracellular accumulation using flow cytometry or ELISA. A) IL-6 levels (pg/ml) in the media were measured using ELISA or C) Flow cytometry following treatment of astrocytes with Aβ oligomers. B) IL-8 levels (pg/ml) in the media were measured using ELISA or D) Flow cytometry (fold change) following treatment of astrocytes with Aβ oligomers. E) Secreted levels of GFAP in the media or F) Cell lysates were measured using ELISA (ng/ml). Results are expressed as ± SD, n=3 P<0.05 (*), P<0.01 (**), P<0.001 (***). Comparisons between treatments were performed using ANOVA followed by Dunnet’s post-test.

### Altered AβPP processing in astrocytes carrying PSEN1 mutations

Whilst treatment with Aβ provides some insight into the impact of acute amyloid exposure, fAD iPSC derived astrocytes may provide a more physiological assessment of chronic APP misprocessing and exposure to Aβ. Although PSEN1 mutations are typically associated with a shift towards longer Aβ isoform accumulation, there is evidence of differences between mutation sites ^50^. Thus, ‘pooled’ PSEN1 lines and individual lines were compared to controls. To quantify AβPP processing in fAD iPSC derived astrocytes carrying PSEN1 mutations compared to ‘healthy’ control, markers associated with amyloidogenic and non-amyloidogenic processing such as Aβ1-40 and Aβ1-42 levels were measured. As expected, Aβ1-40 was significantly elevated in pooled PSEN1 astrocytes compared to control astrocytes (14.67 ± 2.230 vs.11.58 ± 0.806 pg/mg, *p* = 0.055) as quantified by ELISA. Analysis of iPSC derived astrocytes carrying a R278I mutation (15.16 ± 0.543 pg/mg, *p* = 0.012) and A246E (16.64 ± 1.042 pg/mg, *p* = 0.002) mutations demonstrated significantly elevated Aβ1- 40 compared to control astrocytes, however there was no detectable difference in the L286V mutation (12.20 ± 1.778 pg/mg, *p* > 0.05) (Figure 4-D). Aβ1-42 was significantly elevated in PSEN1 astrocytes compared to control (4.283 ± 0.892 vs. 1.868 ± 0.810 pg/mg, *p* = 0.002). This was observed in all PSEN1 mutations, R278I (5.342 ± 0.672 pg/mg, *p* = 0.002), L286V (3.861 ± 0.319 pg/mg, *p* = 0.007) and A246E (3.645 ± 0.266 pg/mg, *p* = 0.012) compared to control (Figure 4-E). In AD, the ratio Aβ1-42:40 is elevated, we next tested whether this was also seen in PSEN1 astrocytes. Taking the ratio of Aβ1-42:40, there was an increase in PSEN1 astrocytes compared to control (0.299 ± 0.074 vs. 0.1607 ± 0.064 pg/mg, *p* = 0.017). Interestingly, this ratio was not elevated in the A246E mutation, however, aggregated-Aβ was significantly increased at baseline in A246E (24.51 ± 4.461 ng/mL, *p* = 0.02) astrocytes only, compared to control (11.07 ± 1.781 ng/mg) (Figure 4-F&G). Together, this suggests that fAD-associated mutations to PSEN1 in astrocytes affect APP processing and therefore will enable us to determine the impact of altered amyloid production of metabolic and disease associated phenotypes.

**Figure 4.**
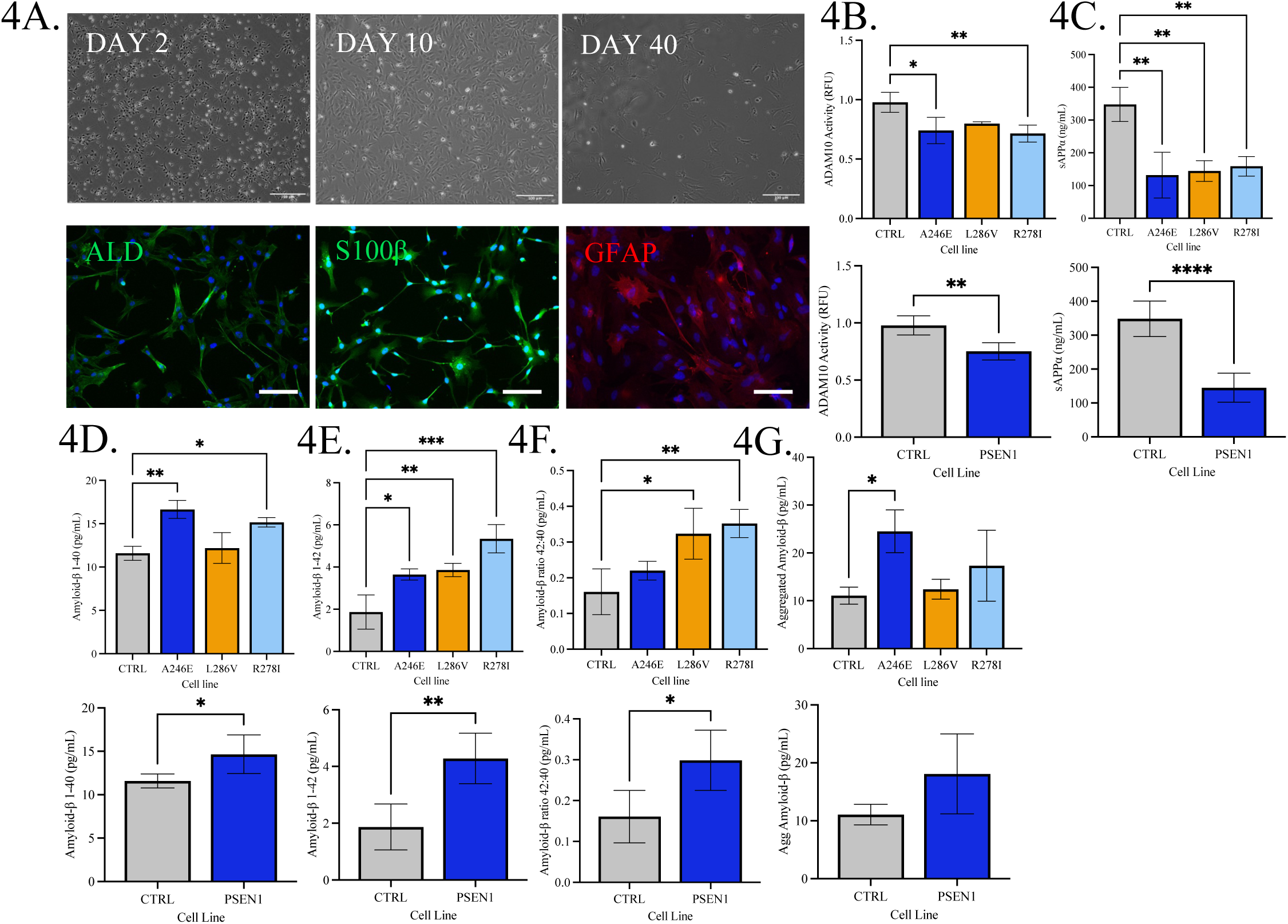
Astrocytes process APP differentially in ‘healthy’ control vs fAD patient derived cells. Control and fAD (PSEN1 mutation) derived astrocytes were characterised using immunofluorescence and APP processing was assessed using ELISA and ADAM 10 activity. A) representative images of fAD (PSEN1 mutation) derived astrocytes. Cells were stained using immunofluorescent antibodies for astrocytic markers ALDH1A1 (green), S100β (Green) and GFAP (red). nuclei were counterstained with DAPI (blue). Scale bars: 100μM. B) Characterisation of ADAM10 enzymatic activity (RFU) and soluble APPα (ng/ml) in control and PSEN1 (A246E, L268V and R278I). Pooled PSEN1 samples compared to control are displayed. D) Aβ 1-40 (pg/ml) Aβ 1-42 (pg/ml), F) Aβ42/40 ratio, G) Aggregated Aβ (pg/ml) were measure in control and fAD patient derived astrocytes after 48 hours. Pooled PSEN1 samples compared to control are displayed. Results are expressed as ± SD, n=3 P<0.05 (*), P<0.01 (**), P<0.001 (***). For direct comparison between control and PSEN1, unpaired t-tests were performed. Comparisons between individual PSEN1 lines were performed using ANOVA followed by Dunnet’s post-test.

To understand how non-amyloidogenic AβPP processing is affected in astrocytes carrying PSEN1 mutations, ADAM10 activity and sAβPPα were quantified. ADAM10 activity was significantly lower in PSEN1 astrocytes compared to control (0.978 ± 0.084 vs. 0.751 ± 0.076 RFU *p* = 0.001). This was significantly lower in R278I (0.715 ± 0.071 RFU, *p* = 0.009) and A246E (0.741 ± 0.112 RFU, *p* = 0.02) mutations compared to control. Additionally, sAβPPα was significantly lower in R278I (158.7 ± 29.64 pg/mg, *p* = 0.004), L286V (144.5 ± 31.63 pg/mg, *p* = 0.002) and A246E (131.7 ± 70.00 pg/mg, *p* = 0.002) compared to control (347.9 ± 52.15 pg/mg) (Figure 4-A&B).

### PSEN1 carrying astrocytes show features of altered metabolism and glutamate uptake

As treatment with Aβ1-42 oligomers resulted in a significant shift in metabolic profile, we investigated the profiles of PSEN1 carrying-astrocytes compared to healthy controls, OCR and ECAR were quantified using Seahorse assay. Basal OCR was elevated in PSEN1 astrocytes compared to control (14.11 ± 1.789 vs. 10.79 ± 1.181 pmol/min, *p* = 0.001), however when looking at individual mutations, this was only altered in R278I (13.88 ± 1.031 pmol/min, *p* = 0.004) and A246E (15.75 ± 0.880 pmol/min, *p* < 0.001) mutations but not in L286V. After correcting for basal OCR, maximal respiratory capacity was significantly elevated in all PSEN1 mutations compared to control (15.12 ± 2.428 vs. 5.580 ± 0.722 pmol/min, *p* < 0.001).

Proton leak (5.065 ± 0.580 vs. 3.260 ± 0.372 pmol/min, *p* = 0.001) and non-mitochondrial oxygen consumption (3.114 ± 0.588 vs. 1.721 ± 0.317 pmol/min, *p* < 0.001) was also significantly elevated in PSEN1 mutations compared to control, whereas ATP linked respiration was only elevated in A246E astrocytes. Not only was OCR elevated, but there was also an increase in ECAR at baseline in PSEN1 mutants compared to controls indicating a greater overall cell energy demand (2.717 ± 0.325 vs. 1.794 ± 0.146 mpH/min, *p* < 0.001). Glycolytic rate, corrected for or basal rates, was also elevated in PSEN1 astrocytes following the addition of oligomycin to astrocytes (1.519 ± 0.266 vs. 1.103 ± 0.132 mpH/min, *p* = 0.001) (Figure 5-A-C).

**Figure 5.**
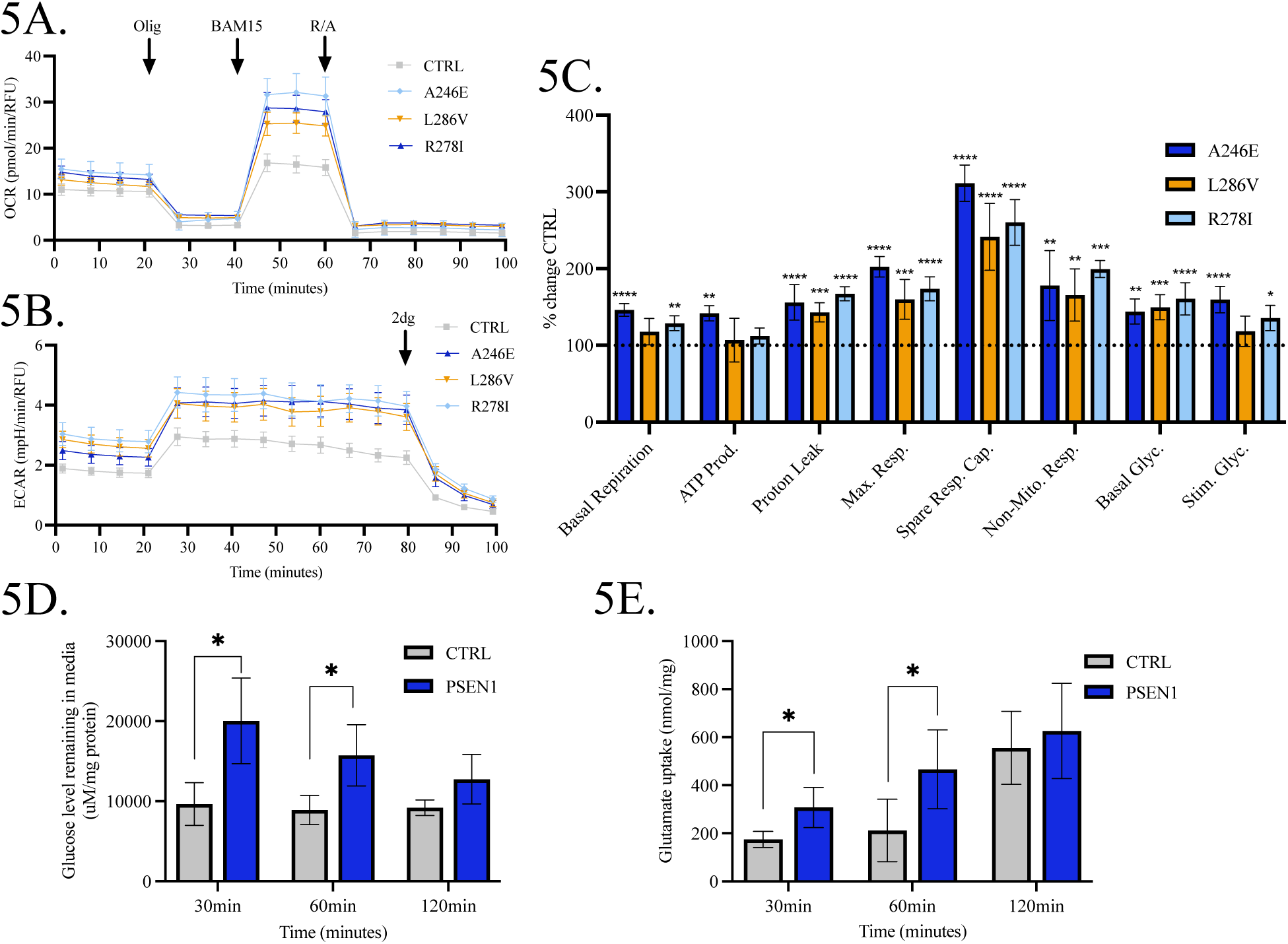
fAD derived astrocytes exhibit differences in their bioenergetic profiles and metabolite processing compared to ‘healthy’ control cells. A) Profiles of Seahorse XFp Mito Stress Test data for oxidative consumption rates (OCR, pmol/min) B) Extracellular acidification rate (ECAR, mpH/min) C) Percentage change of fAD cells over control. D) Glucose levels remaining in the media (µg/mg cellular protein), E) Glutamate uptake (nmol/mg cellular protein) following addition of Glutamate (200µM) over 30-, 60- and 120-min. Results are expressed as ± SD, n=3 P<0.05 (*), P<0.01 (**), P<0.001 (***). Comparisons between treatments were performed using ANOVA followed by Dunnet’s post test 6A.

To understand whether the PSEN1 mutations were associated with functional metabolic processes such as glutamate uptake and lactate release, astrocytes were incubated with media containing 200mM glutamate and sampled at 30 minutes, 60 minutes, 120 minutes. Following glutamate treatment PSEN1 astrocytes displayed an elevated uptake of glutamate at 30 minutes (307.7 ± 83.91 vs. 174.2 ± 33.32 µmol/mg, *p* = 0.028) and 60 minutes (466.3 ± 164.5 vs. 212.5 ± 130.0 µmol/mg, *p* = 0.038) post treatment, however, this was not significantly different to control at 120 minutes. Glutamate remaining in cell culture media after 48hrs was significantly elevated in PSEN1 astrocytes compared to control (2723 ± 345.5 vs. 1968 ± 222.6 µmol/mg, *p* = 0.006). Glutamate uptake can be a highly energy consuming process due to the additional ATP demands of exchanging Na^+^ and K^+^ ions. Thus, glucose uptake from cell culture media and lactate release were measured. Interestingly, despite elevated glutamate uptake there was no significant change in lactate release for any time points, however, glucose remaining in cell culture media was elevated at 30 minutes (20048 ± 5349 vs. 9648 ± 2661 µmol/mg, *p* = 0.010) and 60 minutes (15729 ± 3828 vs. 8912 ± 1814 µmol/mg, *p* = 0.016) post glutamate treatment. There was also significantly more glucose remaining in cell culture media at 48hrs in PSEN1 astrocytes compared to controls (11388 ± 2049 vs. 6644 ± 2756 µmol/mg, *p* = 0.009) (Figure 5-D&E)

### PSEN1 carrying astrocytes display features of reactive gliosis and oxidative stress

A number of studies have described reactive astrocytes in postmortem brain of patients with AD and pre-symptomatically in mouse models of AD suggesting that astrocytes are reactive in AD. As with measurements in Aβ1-42-treated astrocytes, markers associated with a reactive astrocytic state were quantified in PSEN1 mutation cells. Flow cytometric analysis of cytokine accumulation under resting conditions were quantified following protein trafficking inhibition. Both IL-6 (16.29 ± 2.243 vs. 1.028 ± 0.120 FC, *p* < 0.001) and IL-8 (10.19 ± 2.663 vs. 1.036 ± 0.140 FC, *p* = 0.002) were significantly higher in PSEN1 astrocytes than controls respectively (Figure 6 D&E). Further, measurements of GFAP secreted into cell culture media was significantly elevated in PSEN1 astrocytes (3.523 ± 0.468 vs. 2.870 ± 0.028 ng.mg, *p* =0.041), this was also seen in cell lysates compared to controls (23.50 ± 3.249 vs. 18.25 ± 0.851 ng/mg, *p* = 0.023). Mitochondrial superoxide generation, measured via Mitosox Red was not significantly different between PSEN1 and control when grouped, although, one of the mutations, A246E, was significantly elevated compared to control (309400 ± 23669 vs. 223455 ± 17610 RFU, *p* = 0.0009) in specific PSEN1 mutation analysis. However, lipid oxidation measured as total F2 8-isoprostane was significantly elevated in all PSEN1 mutant lines as compared to control (5.510 ± 1.919 vs. 0.5907 ± 0.4420 pg/mg, *p* = 0.0016) (Figure 6-A-C).

**Figure 6.**
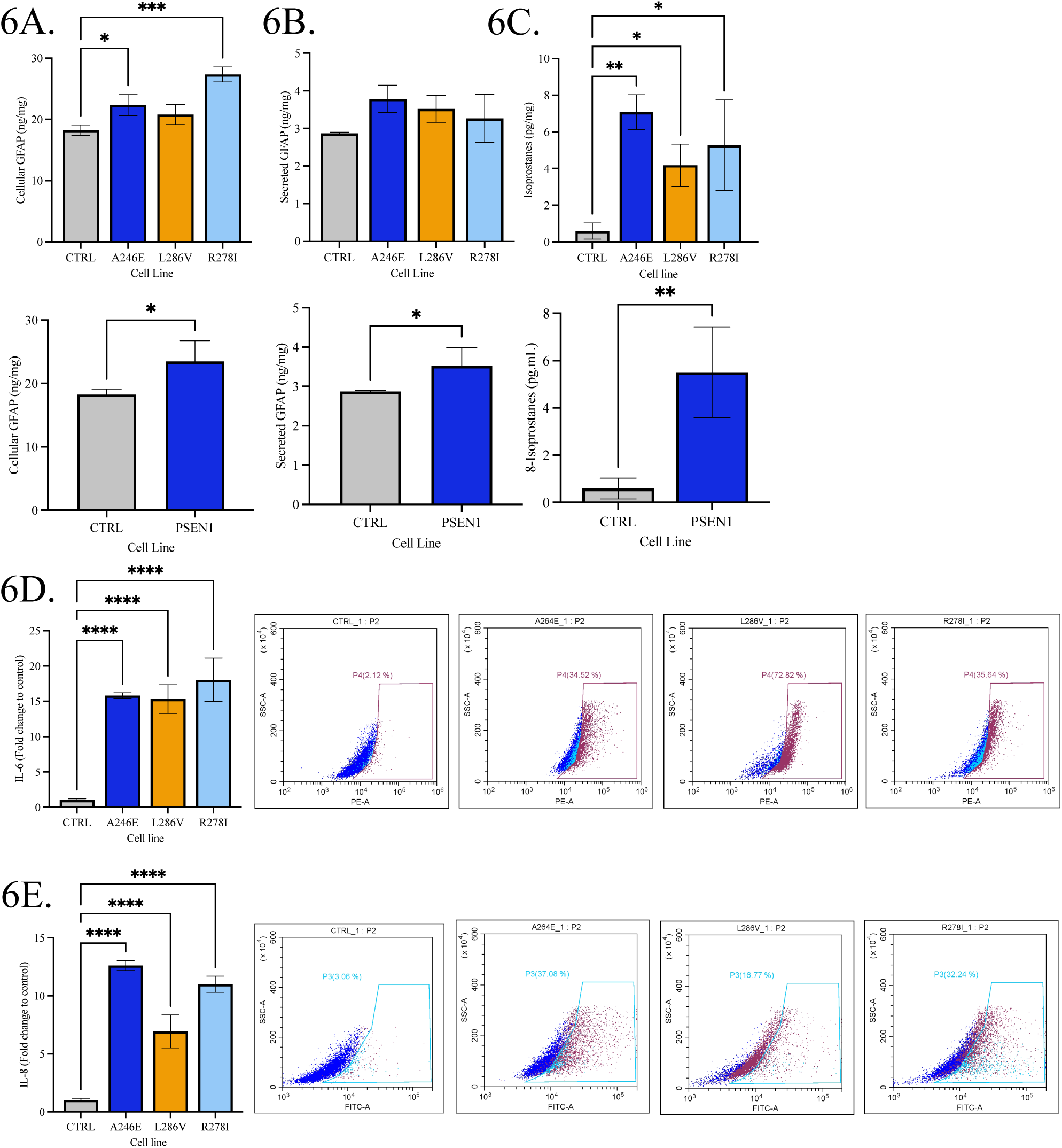
fAD Astrocytes display markers of gliosis compared to ‘Healthy’ control astrocytes. Cytokine, GFAP, 8-Isoprostane levels in media or cellular lysates were measured in control and fAD astrocytes (45 days old). A) Levels of GFAP were measure in the cell lysates or F) cell culture media using ELISA (ng/ml). C) Isoprostane levels were also measured in cellular lysates (pg/mg). Pooled control and fAD cell samples are compared. D) IL-6 and E) IL-8 levels were measured using Flow cytometry (fold change) following in control and fAD astrocytes. Results are expressed as ± SD, n=3 P<0.05 (*), P<0.01 (**), P<0.001 (***). For direct comparison between control and PSEN1, unpaired t-tests were performed. Comparisons between individual PSEN1 lines were performed using ANOVA followed by Dunnet’s post-test.

### Astrocytes carrying fAD-associated mutations present an altered metabolome

As metabolites including glucose, glutamate and lactate demonstrated altered release and uptake, metabolite profiling of astrocyte CM from cells carrying fAD mutation and controls was performed by the application of high-resolution mass spectrometry. Based on the results of univariate analysis, metabolites with adjusted *p* < 0.05 and fold change > 2 were selected as significant. The heatmap plots demonstrate a significantly altered abundance in metabolites in PSEN1 astrocytes compared to control, with L286V astrocytes displaying an altered metabolism to both A246E and R278I astrocytes (Figure 7-A). For multivariate analysis, the PCA models were built to evaluate the similarities among the variables and the robustness of analytical system. The pooled QC samples were clustered tightly towards the centre of the scores plot, indicating that the satisfactory stability of instrument was achieved. Clear clustering and separation for different mutant groups and control groups were observed, suggesting that astrocytes carrying-fAD mutations present a significantly different metabolic profiles as compared to controls (Supplementary figure 2-A-D).

**Figure 7.**
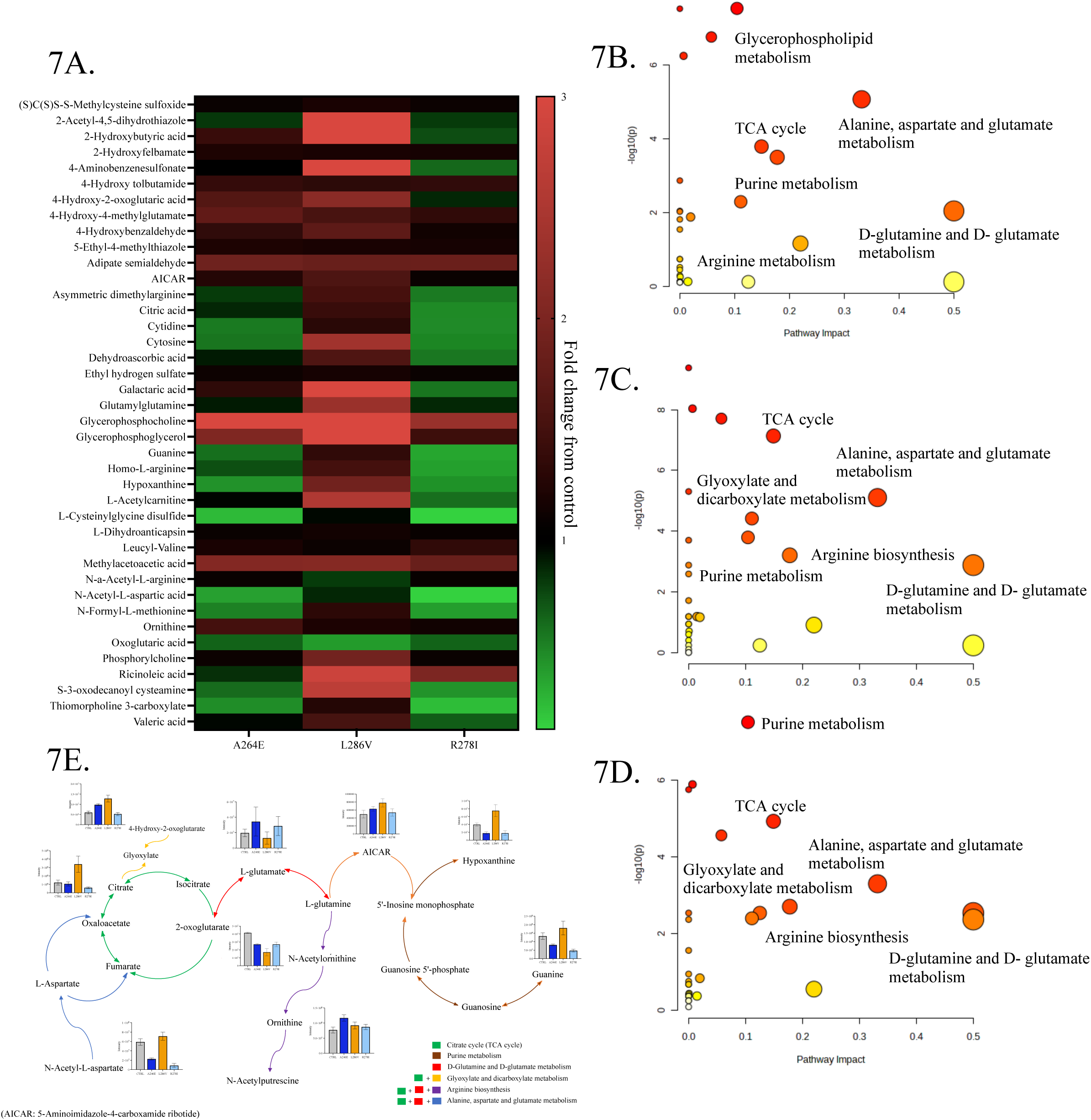
fAD derived astrocytes display significantly altered metabolic profiles compared to controls. A) Heatmap displaying differential compounds identified using metabolomic analysis. Summary of Pathway analysis for the comparison of control and B) A246E, C) L286V and D) R278 fAD patient derived astrocytes. The pathway analysis results of PSEN1 astrocytes compared with control: the colour graduated from white to and red indicates the degree of significance, the size of bubble represents the number of metabolites hit in the pathway. E) Bar charts indicating intensity changes in key metabolites represented within metabolic pathway that were detected in astrocytes carrying PSEN1 mutations. Data shown is expressed as ± SD, n=6.

**Figure 8.**
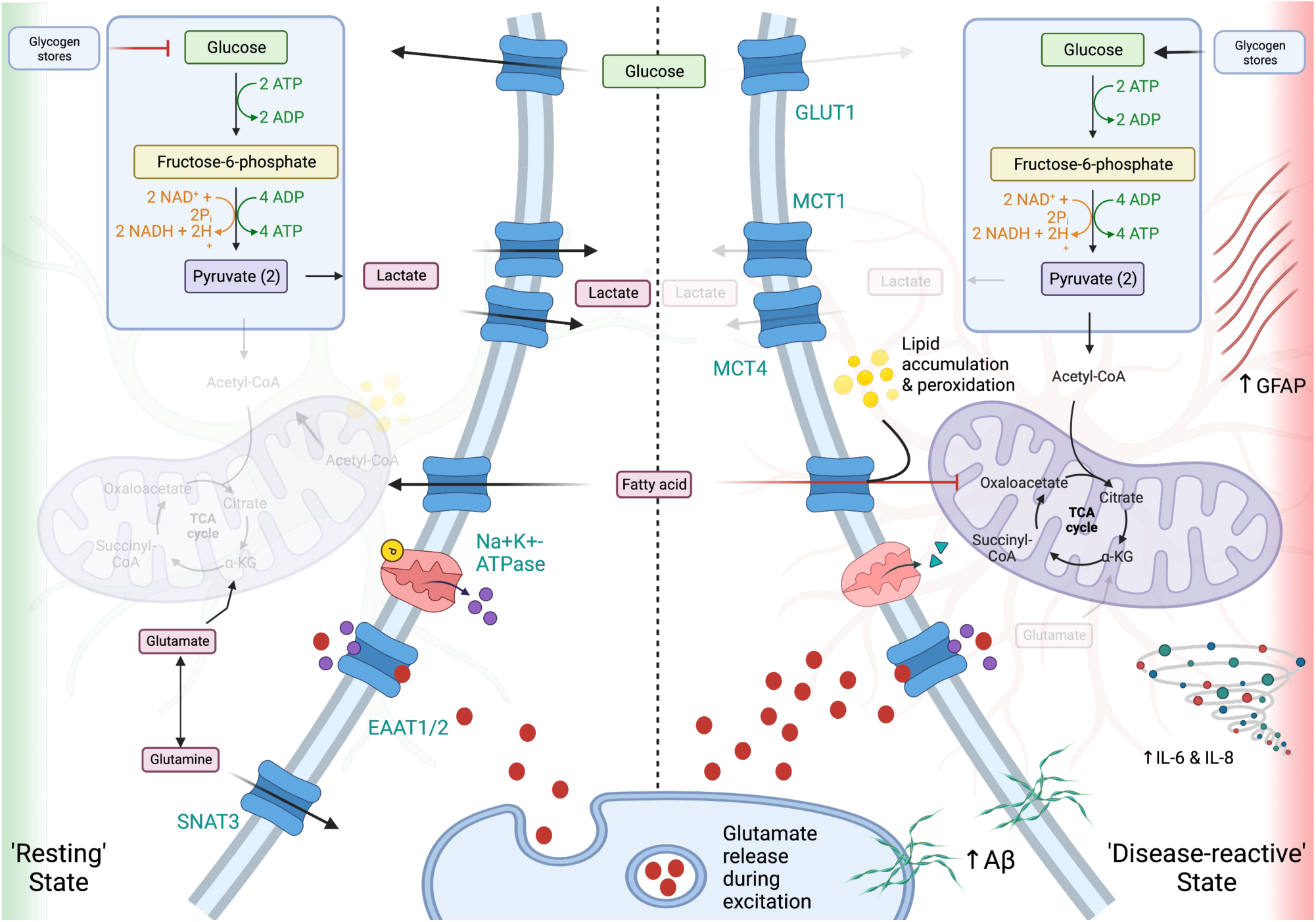
Proposed key metabolic and biochemical changes effecting astrocytes under resting conditions and in a ‘disease-reactive state’ induced by Aβ oligomers or in astrocytes carrying PSEN1 mutations. Astrocytes exposed to Aβ oligomers following exogenous treatment or through endogenous production in PSEN1 mutation carrying cells display and altered metabolism. Under resting conditions, glutamate uptake from excitatory neurons from the synapse is regulated however, this is altered with increase Aβ initially increasing uptake, over a short period. Glutamate uptake is an energy costly process, astrocytes are unable to take up glucose efficiently and rely on breakdown of glycogen stores to fuel metabolic activity, this is coupled with a shift towards oxidative phosphorylation thus, lactate release is reduced. However, this is not coupled efficiently to ATP production. Increased lipids accumulate in astrocytes due to reduced transport into the mitochondria and lipid peroxidation is increased. Astrocytes display elevated cytokines and increase GFAP indicative of a disease-reactive state.

The OPLS-DA models were next constructed to identify the differential compounds between the PSEN1 and control groups. On the OPLS-DA scores plot, mutation and control groups were clearly clustered and separated, indicating the significant distinction of metabolic profile between each PSEN1 type and control astrocytes. Combined with cross-validation results and permutation results, the models constructed for this study were less likely to overfit our dataset, with good fitness and good predictive ability (R^2^Y > 0.9 and Q^2^> 0.9). The lists of variables importance in projection (VIP) were then generated for each comparison. VIP value generally represents the importance of variables in the OPLS-DA model and has been used to extract the metabolites related to separation ^51^ (Supplementary figure 2-A-D). Finally, in this study, the metabolites with VIP>1 and adjusted *p*<0.05 were considered as key compounds. As compared to control astrocytes, 8, 8 and 10 differential metabolites (Level 1 and Level 2) were significantly dysregulated in astrocytes carrying A246E, L286V, R278I, respectively.

To identify which metabolic pathways are dysregulated in AD astrocytes, pathway analysis was performed using the all the identified metabolites. Purine metabolism, alanine, aspartate and glutamate metabolism, and citrate (TCA cycle) metabolism were the three most dysregulated pathways for astrocytes carrying A246E and R278I mutations. The metabolites mapped onto these pathways were significantly less abundant in astrocytes carrying A246E and R278I mutations than in control astrocytes. For astrocytes carrying the L286V mutation, different from A246E and R278I mutations, the metabolites involved purine metabolism, alanine, aspartate and glutamate metabolism, and TCA cycle were more abundant in L286V mutated astrocytes compared with control astrocytes. Due to a higher impact score on the pathway analysis, the D-glutamine and D-glutamate metabolism was selected to compare L286V mutations with control, with metabolites in this pathway (including L-glutamate and 2- oxoglutarate) significantly reduced as compared to control astrocytes. Together, the data presented demonstrates that astrocytes carrying AD mutations present a dysregulated metabolism and that different AD mutations can have different effects on astrocyte metabolism (Figure 7-B-E).

## Discussion

Astrocytes are key mediators in cerebral metabolism and may even carry a similar metabolic cost to neurons which are typically thought of as the most energetically demanding cells in the brain ^52,53^. Disruption to astrocytic function, either caused by, or resulting in, a reduction in glucose uptake, may result in a decline in homeostatic control, thereby reducing the ability of cells to respond to neuronal activity and stress. In order to be able to utilise iPSC-derived cells to study the early metabolic changes in AD, it is necessary to first ensure that the astrocytes behave similarly to cells ‘*in vivo’*. We and others have shown that astrocytes differentiated from iPSCs show typical stellate morphology, and express astrocyte markers including S100β, ALDH1 and GFAP ^29,54,55^. In the brain astrocytes breakdown glycogen under conditions of starvation in order to maintain survival of neurons, protect axons, and ensure synaptic activity is maintained ^22^. Furthermore, during neuronal activity uptake of glutamate and NaCl^2^ by glutamate transporters in astrocytes initiates glycogenolysis and rapid lactate production ^56^, which may be used as a fuel source in neurons (astrocyte-neuron lactate shuttle), although this is still a debated area ^57–59^. Data presented here shows that glutamate induces glycogenolysis in astrocytes and that process can be blocked using Ouabain and TBOA demonstrating that glutamate uptake and efflux of NaCl^2+^ is associated with glycogen breakdown. Furthermore, we demonstrate that astrocytes have the ability to store glycogen and are able to undergo glycogenolysis in response to hypoglycaemia and in response to dbcAMP. Whilst these findings have been reported in primary rodent cells ^60–63^ and human NT2.D1 embryo carcinoma derived astrocytic cells ^64^, this is the first time that this response has been reported in human iPSC derived astrocytes.

Evidence of altered metabolic profiles in iPSC-derived astrocytes from late-onset AD has been previously shown ^65^. Aβ is an important hallmark of AD pathology and is postulated to play a role in altered metabolism, therefore, we investigated how relatively acute treatment of Aβ altered astrocytic metabolism. iPSC astrocytes showed a concentration-dependent pattern of reduction in glucose uptake which concurs with previous work^38,66^. The pathogenesis of amyloid-induced reduction in glucose uptake remains a focus of research, and several mechanisms have been suggested as possible pathways for glucose uptake reduction. For example, Aβ reduces glucose uptake by preventing GLUT3 fusion to the plasma membrane, despite elevated protein expression ^67^. Impaired lipid peroxidation has also been proposed as a mechanism for reduced glucose uptake in AD ^68^. Further, the intracellular aggregation of Aβ can induce alterations in pro-inflammatory and metabolic reprogramming that predisposes cells to die through regulated cell death pathways, a process which can induce change in glucose metabolism and mitochondrial bioenergetics ^69^. In this study, treatment with Aβ1-42 oligomers induced a significant reduction in glycogen content. The mechanisms behind this amyloid-induced reduction in glycogen turnover remain to be explained but is perhaps unsurprising since cells exhibit a reduction in glucose uptake and may therefore initiate glycogenolysis. In addition, we observed alterations in basal metabolic rates (increase oxygen consumption) and differences in glutamate uptake and lactate release following acute Aβ treatments. This data shows that Aβ may interfere with glutamate uptake, which would impact one of the key roles of astrocytes in mediating neurotoxicity. This in turn would hinder lactate shuttling which is a key energy source for neurons provided by astrocytes during excitation or low glucose availability. However, further research utilising the co-culture of neurons and astrocytes is needed to confirm dysfunction of cell-cell metabolic coupling. We demonstrate that iPSC-derived astrocytes display features associated with a reactive state following Aβ exposure, which is in agreement with previous studies that show AD pathology in PSEN1 astrocytes ^29^. In fact, similar findings are demonstrated here, with increased Aβ generation and an increased Aβ42:40 ratio in PSEN1 astrocytes. We add further evidence of a reduction in non-amyloidogenic AβPP processing via lower ADAM10 activity and reduced sAβPPα secretion was detected in PSEN1 astrocytes compared to controls. The relationship between the presence of the PSEN1 mutation and reduced ADAM10 activity in astrocytes is not known, and remains an interesting topic for further investigation, ^70^.

It was hypothesised PSEN1 astrocytes which are chronically exposed to elevated, (albeit much lower) levels of Aβ, would display metabolic alterations and reactivity. Under resting conditions, we detected significantly higher oxygen consumption compared to control astrocytes, which has been previously shown^29,65^. Interestingly, we also demonstrate an elevation in non-mitochondrial oxygen consumption, proton leak and a reduction in ATP-linked oxygen consumption suggesting an alteration in the efficiency of the astrocytic aerobic system and early hypermetabolic phenotype even with a reduction in glucose uptake. This elevated metabolic profile has been previously reported in late onset AD derived cells ^65^. Further to this, a marked elevation in maximal uncoupled aerobic capacity was detected, suggesting an adaptive response to cellular stress in the PSEN1 mutation and highlights areas for future work. PSEN1 astrocytes used in this study also demonstrated a pronounced reactive state, indicated by increased cell size, elevated inflammatory cytokine accumulation and an increase in GFAP protein levels. This supports the growing evidence of a role of reactive astrocytes in mediating aspects of AD pathology and may be a key feature of early pathogenesis ^27,71,72^. Furthermore, evidence of redox stress was identified with significantly increased lipid peroxidation, although this was not matched by any alteration in superoxide generation. The majority of lipid peroxides have been attributed to neuronal sources although evidence of astrocytic lipid peroxidation has been previously shown in *in vivo* AD models ^73,74^. Metabolomic analysis of patient derived and Aβ treated astrocytes revealed significant alterations in key pathways including citric acid cycle, purine metabolism, glutamine, glutamate, arginine, alanine and aspartate metabolism. Alterations in key metabolic pathways have previously been reported in AD ^75–78^.

Studying the early metabolic features of AD is crucial for developing effective treatments and understanding the disease process. Data presented herein shows altered markers of AD pathology, including APP misprocessing and Aβ production, in fAD patient derived astrocytes. Furthermore, we demonstrate significant metabolic changes in both Aβ treated and fAD derived astrocytes, as well as elevated markers of astrocyte reactivity. Both fAD and Aβ treated astrocytes demonstrate a bioenergetic shift to a hypermetabolic state despite a reduced uptake of glucose and glutamate. Our results highlight the impact of AD on the anaplerotic and cataplerotic nature of astrocytes and demonstrate how the delicate balance of maintaining neurotransmitter and metabolic demands could be perturbed in response to APP misprocessing in both acute and chronic exposure to Aβ. These findings emphasize the importance of astrocytes in cerebral metabolism and highlight how this could be perturbed in neurodegeneration. Treatments targeting astrocytic reactivity as well as metabolic dysfunction may reduce or ameliorate the further development of AD and should be considered in preventative early trials in preclinical studies.

## Acknowledgments

These cell lines were produced from fibroblasts that were kindly donated by people from families with genetic forms of dementia as part of the study ‘Induced pluripotent stem cells derived from patients with Familial Alzheimer’s disease and other dementias as novel cell models for neurodegeneration study’ (Ref: 09/0272) under the supervision of Professor Nick Fox at the UCL Queen Square Institute of Neurology Dementia Research Centre. This work was supported by the National Institute for Health and Care Research University College London Hospitals Biomedical Research Centre. For further information about the cell lines please contact Professor Selina Wray at.

