## Supplemental figures for "Altered metabolic function induced by amyloid-beta oligomers and PSEN1-mutations in iPSC-derived astrocytes"

### For supplementary

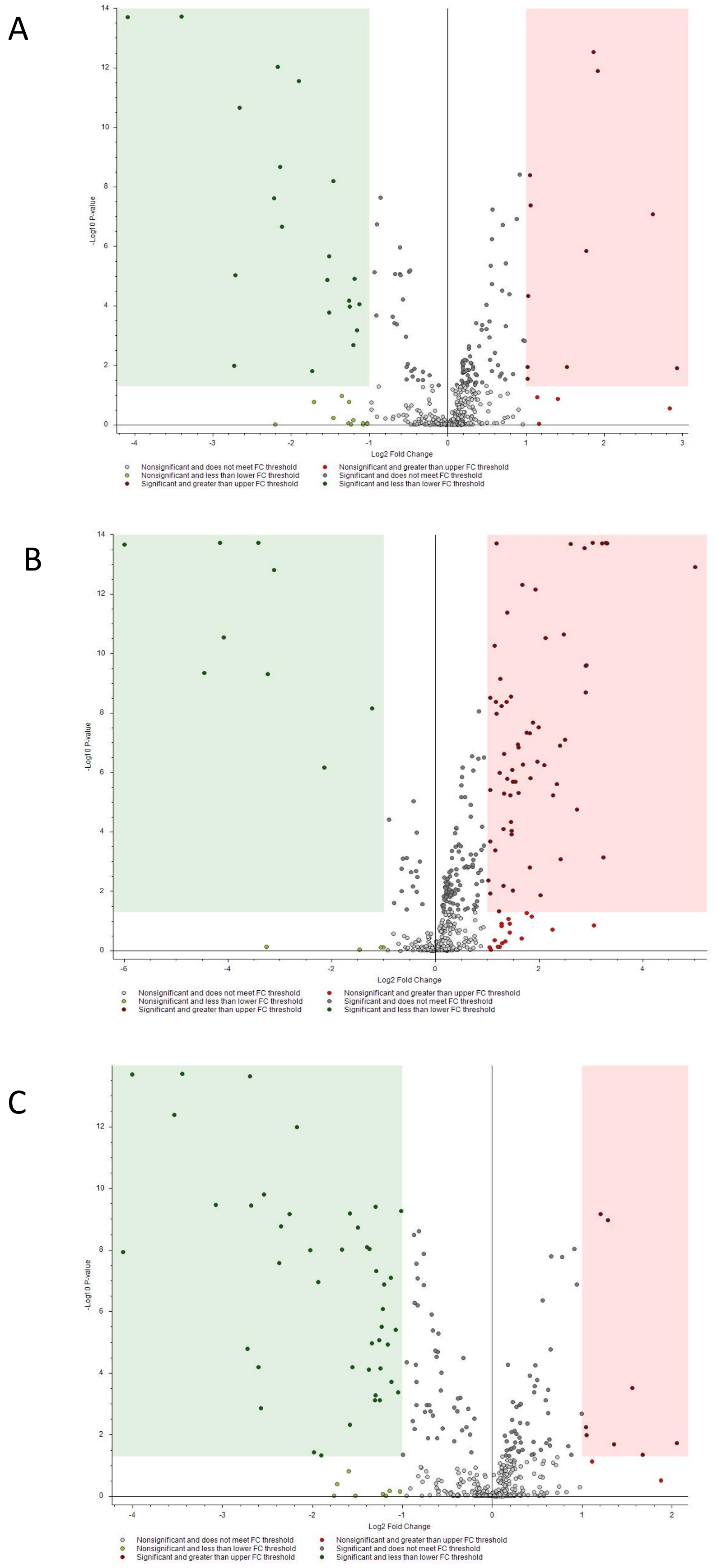


Sup. Figure 1. The volcano plots of PSEN1 astrocytes compared with control: metabolites in red were more abundant in PSEN1 astrocytes while metabolites in green were identified at higher level in control relative to PSEN1 astrocytes. (A): A246E vs. control. (B): L286V vs. control. (C): R278I vs. control.

**For supplementary**

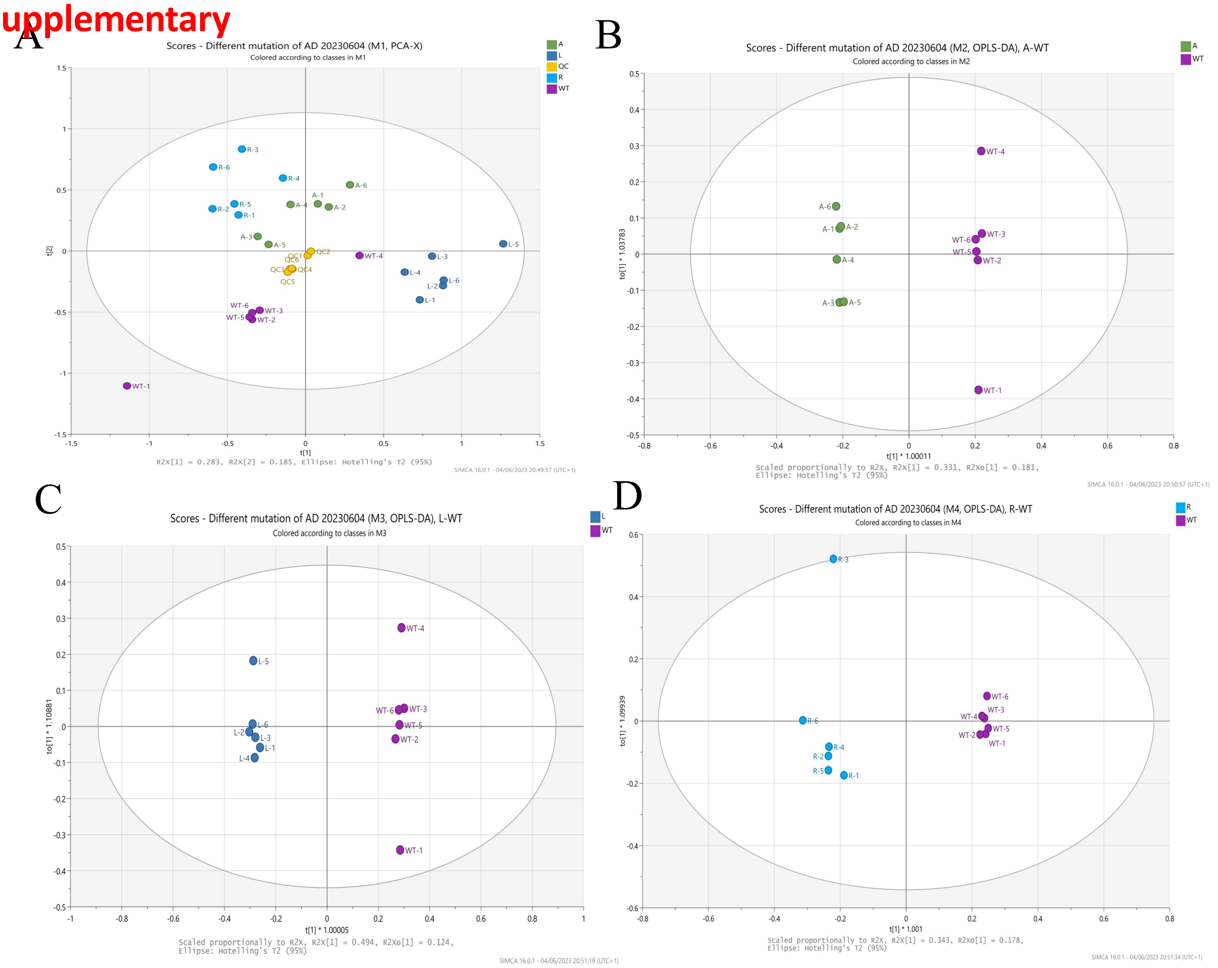


Sup. Figure 2. Summary of results of PCA and OPLS-DA results for the comparison of PSEN1 mutations with control astrocytes. (A): The PCA scores plot of different PSEN1 mutation astrocytes and control astrocytes. (B): The OPLS-DA scores plot of A264E (A) compared with control (WT): $R^2Y$ 0.999 and $Q^2$ 0.939. (C):The OPLS-DA scores plot of L286V compared with control: $R^2Y$ 0.998 and $Q^2$ 0.967. (D):The OPLS-DA scores plot of R278I mutant compared with wild type: $R^2Y$ 0.987 and $Q^2$ 0.931.

**For supplementary**

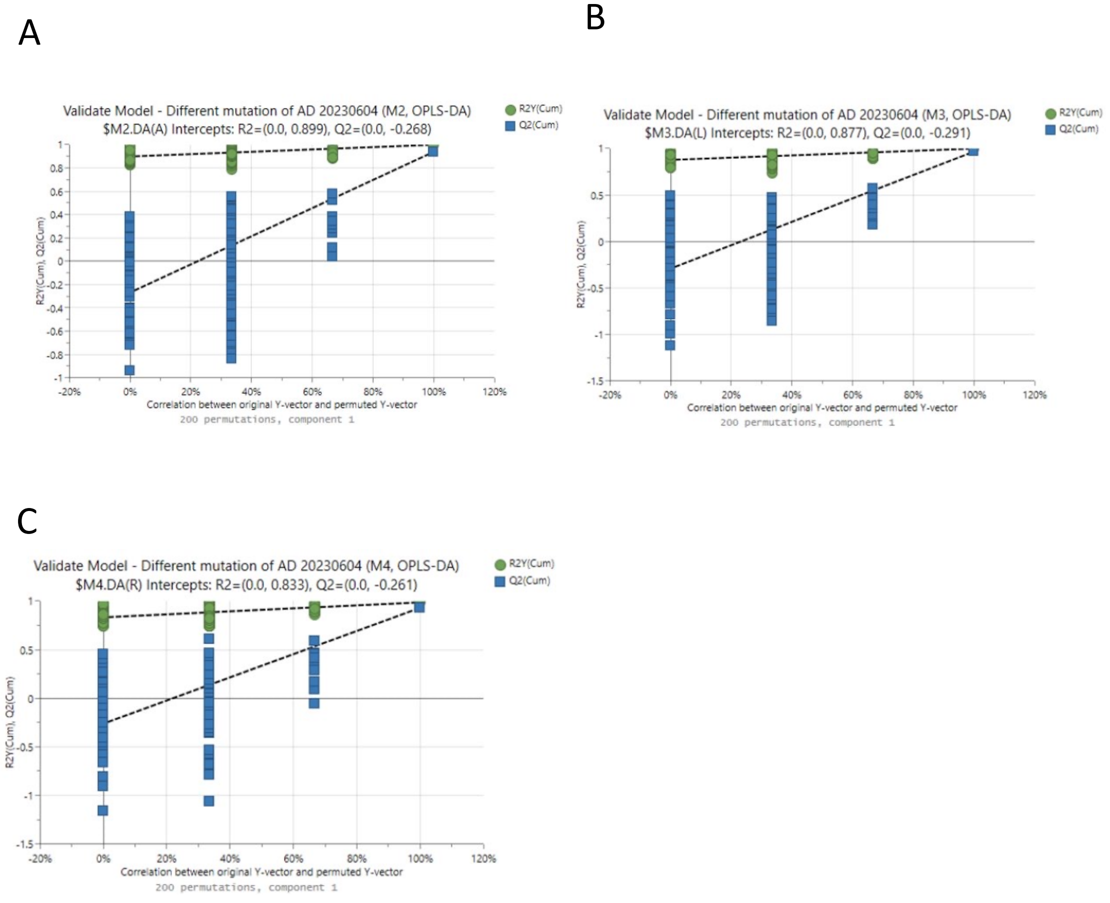


Sup. Figure 3. The permutation test results of each comparison. (A): The permutation test of A264E mutation compared with control. (B): The permutation test of L286V mutation compared with control. (C): The permutation test of R278I mutant compared with control.

**Table 1. The results of significant pathways for A264E mutant compared with control (adjusted p< 0.05 and impact factor >0.1).**

| Pathway name | Total Compounds | Hits | Raw p | Holm adjust | FDR | Impact |
|---|---|---|---|---|---|---|
| Purine metabolism | 65 | 3 | 2.87E-08 | 8.62E-07 | 4.40E-07 | 0.10424 |
| Alanine, aspartate and glutamate metabolism | 28 | 4 | 8.36E-06 | 2.17E-04 | 5.02E-05 | 0.33174 |
| Citrate cycle (TCA cycle) | 20 | 2 | 1.60E-04 | 3.99E-03 | 7.98E-04 | 0.14894 |
| Arginine biosynthesis | 14 | 3 | 3.12E-04 | 7.49E-03 | 1.34E-03 | 0.17766 |

**Table 2. The results of significant pathways for L286V mutant compared with control (adjusted p< 0.05 and impact factor >0.1).**

| Pathway name | Total Compounds | Hits | Raw p | Holm adjust | FDR | Impact |
|---|---|---|---|---|---|---|
| Citrate cycle (TCA cycle) | 20 | 2 | 7.35E-08 | 1.99E-06 | 5.51E-07 | 0.14894 |
| Alanine, aspartate and glutamate metabolism | 28 | 4 | 7.81E-06 | 1.95E-04 | 3.90E-05 | 0.33174 |
| Glyoxylate and dicarboxylate metabolism | 32 | 3 | 3.82E-05 | 9.17E-04 | 1.64E-04 | 0.11112 |
| Purine metabolism | 65 | 3 | 1.60E-04 | 3.68E-03 | 5.99E-04 | 0.10424 |
| Arginine biosynthesis | 14 | 3 | 6.20E-04 | 1.30E-02 | 1.86E-03 | 0.17766 |
| D-Glutamine and D-glutamate metabolism | 6 | 2 | 1.30E-03 | 2.61E-02 | 3.26E-03 | 0.50000 |

**Table 3. The results of significant pathways for R278I mutant compared with control (adjusted p< 0.05 and impact factor >0.1).**

| Pathway name | Total Compounds | Hits | Raw p | Holm adjust | FDR | Impact |
|---|---|---|---|---|---|---|
| Purine metabolism | 65 | 3 | 3.11E-08 | 9.32E-07 | 9.32E-07 | 0.10424 |
| Citrate cycle (TCA cycle) | 20 | 2 | 1.18E-05 | 3.19E-04 | 8.85E-05 | 0.14894 |
| Alanine, aspartate and glutamate metabolism | 28 | 4 | 4.99E-04 | 1.25E-02 | 2.50E-03 | 0.33174 |
| Arginine biosynthesis | 14 | 3 | 1.97E-03 | 4.73E-02 | 8.45E-03 | 0.17766 |

**Table 1. The differential metabolites between A264E mutant astrocytes and control astrocytes (Adjusted p<0.05 and VIP >1).**

| m/z | Retention time (min) | Formula | ΔMass [ppm] | Putative Name | HMDB ID | Identification confidence | Reference Ion | Log2 Fold Change | Adjusted P | VIP |
|---|---|---|---|---|---|---|---|---|---|---|
| 258.1101 | 10.03 | $C_8H_{20}NO_6P$ | 0.03 | sn-Glycero-3-Phosphocholine | HMDB0000086 | *m/z*, RT, MS/MS (Level 1) | [M+H]+1 | 1.87 | 1.85E-10 | 2.21516 |
| 133.0971 | 17.06 | $C_5H_{12}N_2O_2$ | -0.17 | Ornithine | HMDB0000214 | *m/z*, RT, MS/MS (Level 1) | [M+H]+1 | 0.56 | 6.31E-04 | 1.24816 |
| 115.0402 | 4.43 | $C_5H_8O_3$ | 1.03 | Methyl acetoacetate | HMDB0000310 | *m/z*, MS/MS (Level 2) | [M-H]-1 | 1.05 | 5.80E-07 | 1.68004 |
| 137.0458 | 8.08 | $C_5H_4N_4O$ | -0.3 | Hypoxanthine | HMDB0000157 | *m/z*, RT, MS/MS (Level 1) | [M+H]+1 | -1.19 | 4.54E-04 | 1.73659 |
| 152.0566 | 9.98 | $C_5H_5N_5O$ | -0.46 | Guanine | HMDB0000132 | *m/z*, RT, MS/MS (Level 1) | [M+H]+1 | -0.68 | 7.76E-03 | 1.35949 |
| 176.0388 | 4.55 | $C_6H_{11}NO_3S$ | 0.34 | N-Formyl-L-methionine | HMDB0001015 | *m/z*, MS/MS (Level 2) | [M-H]-1 | -0.9 | 1.20E-05 | 1.57238 |
| 129.0559 | 4.03 | $C_6H_{10}O_3$ | 1.32 | 6-Oxohexanoic acid | HMDB0012882 | *m/z*, MS/MS (Level 2) | [M-H]-1 | 0.92 | 5.80E-07 | 1.56017 |
| 145.0144 | 10.54 | $C_5H_6O_5$ | 0.79 | 2-Oxoglutarate | HMDB0000208 | *m/z*, RT, MS/MS (Level 1) | [M-H]-1 | -0.65 | 8.24E-03 | 1.28067 |

(Level 1 identification was that metabolites were matched with accurate masses, retention times and MS/MS fragmentation of authentic standards. Level 2 identification was that metabolites were matched with accurate masses and MS/MS information in online spectral library when lack of standards.)

**Table 2. The differential metabolites between L286V mutant astrocytes and control astrocytes (Adjusted p<0.05 and VIP >1).**

| m/z | Retention time (min) | Formula | ΔMass [ppm] | Putative Name | HMDB ID | Identification confidence | Reference Ion | Log2 Fold Change | Adjusted P | VIP |
|---|---|---|---|---|---|---|---|---|---|---|
| 274.10455 | 10.03 | $C_{10}H_{17}N_3O_6$ | -0.05 | L-Glutamyl-L-glutamine | HMDB0028817 | m/z, MS/MS (Level 2) | [M-H]-1 | 1.18 | 2.23514E-12 | 1.38638 |
| 258.11011 | 10.03 | $C_8H_{20}NO_6P$ | 0.03 | sn-Glycero-3-Phosphocholine | HMDB0000086 | m/z, RT, MS/MS (Level 1) | [M+H]+1 | 3.03 | 2.23514E-12 | 2.19947 |
| 115.04019 | 4.431 | $C_5H_8O_3$ | 1.03 | Methyl acetoacetate | HMDB0000310 | m/z, MS/MS (Level 2) | [M-H]-1 | 1.07 | 1.02579E-07 | 1.30344 |
| 137.04575 | 8.08 | $C_5H_4N_4O$ | -0.3 | Hypoxanthine | HMDB0000157 | m/z, RT, MS/MS (Level 1) | [M+H]+1 | 0.94 | 0.002494024 | 1.14349 |
| 191.01982 | 11.62 | $C_6H_8O_7$ | 0.34 | Citric acid | HMDB0000094 | m/z, RT, MS/MS (Level 1) | [M-H]-1 | 1.46 | 7.98614E-05 | 1.48089 |
| 165.0405 | 8.86 | $C_5H_{10}O_6$ | 0.22 | Arabic acid | HMDB0000539 | m/z, MS/MS (Level 2) | [M-H]-1 | 1.25 | 1.66991E-05 | 1.43568 |
| 129.05589 | 4.03 | $C_6H_{10}O_3$ | 1.32 | 6-Oxohexanoic acid | HMDB0012882 | m/z, MS/MS (Level 2) | [M-H]-1 | 0.85 | 2.4824E-07 | 1.17144 |
| 145.01436 | 10.54 | $C_5H_6O_5$ | 0.79 | 2-Oxoglutarate | HMDB0000208 | m/z, RT, MS/MS (Level 1) | [M-H]-1 | -1.21 | 2.0171E-07 | 1.39994 |

(Level 1 identification was that metabolites were matched with accurate masses, retention times and MS/MS fragmentation of authentic standards. Level 2 identification was that metabolites were matched with accurate masses and MS/MS information in online spectral library when lack of standards.)

**Table 3. The differential metabolites between R278I mutant astrocytes and control astrocytes (Adjusted p<0.05 and VIP >1).**

| m/z | Retention time (min) | Formula | ΔMass [ppm] | Putative Name | HMDB ID | Identification confidence | Reference Ion | Log2 Fold Change | Adjusted P | VIP |
|---|---|---|---|---|---|---|---|---|---|---|
| 101.0609 | 4.49 | $C_5H_{10}O_2$ | 1.39 | Valeric acid | HMDB0000892 | m/z, MS/MS (Level 2) | [M-H]-1 | -0.56 | 1.58E-03 | 1.0173 |
| 258.1101 | 10.03 | $C_8H_{20}NO_6P$ | 0.03 | sn-Glycero-3-Phosphocholine | HMDB0000086 | m/z, RT, MS/MS (Level 1) | [M+H]+1 | 1.21 | 6.43E-08 | 1.53243 |
| 174.0408 | 9.89 | $C_6H_9NO_5$ | 0.04 | N-Acetylaspartic acid | HMDB0000812 | m/z, RT, MS/MS (Level 1) | [M-H]-1 | -2.6 | 1.12E-03 | 2.49373 |
| 115.0402 | 4.43 | $C_5H_8O_3$ | 1.03 | Methyl acetoacetate | HMDB0000310 | m/z, MS/MS (Level 2) | [M-H]-1 | 0.95 | 4.55E-06 | 1.26091 |
| 137.0458 | 8.08 | $C_5H_4N_4O$ | -0.3 | Hypoxanthine | HMDB0000157 | m/z, RT, MS/MS (Level 1) | [M+H]+1 | -1.25 | 1.97E-04 | 1.43215 |
| 152.0566 | 9.98 | $C_5H_5N_5O$ | -0.46 | Guanine | HMDB0000132 | m/z, RT, MS/MS (Level 1) | [M+H]+1 | -1.39 | 4.48E-07 | 1.74261 |
| 176.0388 | 4.55 | $C_6H_{11}NO_3S$ | 0.34 | N-Formyl-L-methionine | HMDB0001015 | m/z, MS/MS (Level 2) | [M-H]-1 | -1.3 | 5.38E-08 | 1.67012 |
| 191.0198 | 11.62 | $C_6H_8O_7$ | 0.34 | Citric acid | HMDB0000094 | m/z, RT, MS/MS (Level 1) | [M-H]-1 | -1.25 | 8.79E-03 | 1.35486 |
| 129.0559 | 4.03 | $C_6H_{10}O_3$ | 1.32 | 6-Oxohexanoic acid | HMDB0012882 | m/z, MS/MS (Level 2) | [M-H]-1 | 0.92 | 4.69E-07 | 1.28763 |
| 145.0144 | 10.54 | $C_5H_6O_5$ | 0.79 | 2-Oxoglutarate | HMDB0000208 | m/z, RT, MS/MS (Level 1) | [M-H]-1 | -0.57 | 5.02E-03 | 1.13648 |

(Level 1 identification was that metabolites were matched with accurate masses, retention times and MS/MS fragmentation of authentic standards. Level 2 identification was that metabolites were matched with accurate masses and MS/MS information in online spectral library when lack of standards.)
